# Multifactorial analysis of terminator performance on heterologous gene expression in Physcomitrella

**DOI:** 10.1101/2023.06.30.547182

**Authors:** Paul Alexander Niederau, Pauline Eglé, Sandro Willig, Juliana Parsons, Sebastian N.W. Hoernstein, Eva L. Decker, Ralf Reski

## Abstract

The production of recombinant proteins for health applications accounts for a large share of the biopharmaceutical market. While many drugs are produced in microbial and mammalian systems, plants gain more attention as expression hosts to produce eukaryotic proteins. In particular the GMP-compliant moss Physcomitrella (*Physcomitrium patens*) has outstanding features such as excellent genetic amenability, reproducible bioreactor cultivation, and humanized protein glycosylation patterns. In this study, we selected and characterized novel terminators for their effects on heterologous gene expression. The Physcomitrella genome contains 53,346 unique 3’UTRs (untranslated regions) of which 7,964 transcripts contain at least one intron. Over 91% of 3’UTRs exhibit more than one polyadenylation site, indicating the prevalence of alternative polyadenylation in Physcomitrella. Out of all 3’UTRs, 14 terminator candidates were selected and characterized via transient Dual Luciferase assays, yielding a collection of endogenous terminators performing equally high as established heterologous terminators CaMV35S, AtHSP90, and NOS. High performing candidates were selected for testing as double terminators which impact reporter levels, dependent on terminator identity and positioning. Testing of 3’UTRs among the different promoters NOS, CaMV35S, and PpActin5 showed an increase of more than 1,000-fold between promoters PpActin5 and NOS, whereas terminators increased reporter levels by less than 10-fold, demonstrating the stronger effect promoters play as compared to terminators. The number of polyadenylation sites as well as polyadenylation signals were found to be major determinants of terminator performance. Our results improve the biotechnology platform Physcomitrella and further our understanding of how terminators influence gene expression in plants in general.

**Key message:** Characterization of Physcomitrella 3’UTRs across different promoters yields endogenous single and double terminators for usage in molecular pharming and indicates promoters and terminators to synergistically control gene expression.

## Introduction

The biotechnology sector is estimated to have an economic potential of around three trillion USD per year in 2030 – 2040, not considering downstream and secondary effects (Chui et al., 2020). A large share is attributed to the biopharmaceutical market and the production of recombinant proteins for health applications. While the most established production platforms are based on bacteria, yeast, insect and mammalian cells; plants are an attractive alternative for producing eukaryotic proteins (Reski et al., 2015; Rybicki, 2020; He et al., 2021; Lobato Gómez et al., 2021).

Plant-based expression systems can operate at lower cost, possess high scalability, produce complex and highly glycosylated proteins, and are free of human pathogens and endotoxins (Twyman et al., 2003; Lomonossoff & D’Aoust, 2016; Capell et al., 2020). The most used plant in molecular pharming is *Nicotiana benthamiana*, which has been successfully glyco-engineered to produce ZMapp, a treatment for Ebola based on an antibody cocktail lacking plant-typical *N*-glycan modifications (Margolin et al., 2018). Other promising plant-produced pharmaceuticals still in clinical trials are virus-like particles (VLPs)-based vaccines such as the hemagglutinin-based influenza VLP (D’Aoust et al., 2010, Ward et al., 2020). A promising alternative plant for producing biopharmaceuticals is the GMP (good manufacturing practise)-compliant moss Physcomitrella (*Physcomitrium patens*) (Reski et al., 2018; Decker & Reski, 2020). Physcomitrella has an excellent genetic amenability due to its haploid dominant phase and its high rate of homologous recombination enabling precise gene targeting (Strepp et al., 1998; Reski, 2018). Plant-specific β1,2-xylosylation, α1,3-fucosylation, β1,3-galactosylation, and formation of Lewis A epitopes on *N*-glycans could be abolished via targeted gene knockouts yielding more humanized and homogenic *N*-glycan patterns (Koprivova et al., 2004; Parsons et al., 2012). Another recent advance was the introduction of enzymes necessary for sialic acid synthesis, activation, and linkage to protein *N*-glycans (Bohlender et al., 2020). The protein α-galactosidase A (aGal) produced in Physcomitrella bioreactors (https://www.elevabiologics.com/), intended to treat Morbus Fabry, successfully passed clinical phase I (Shen et al., 2016; Hennermann et al., 2019). Other moss-based products in the pre-clinical phase are human factor H (FH) and synthetic FH-related multitarget proteins MFHR1 and MFHR13. These glycoproteins act as regulators of the human complement system and are potential biopharmaceuticals for treatment of complement-related disorders (Büttner-Mainik et al., 2011; Michelfelder et al., 2017, 2018; Top et al., 2019; Ruiz-Molina et al., 2022a).

A critical factor for choosing a biotechnology production platform for a drug candidate is product yield. To increase yields, bioprocess design is used at different steps during the production workflow. Upstream processes typically deal with genetic engineering of cell lines, bioreactor growth kinetics, media compositions, etc., whereas downstream processes focus on harvest, purification, and formulation of the final product. In Physcomitrella, various efforts were made to optimize media compositions (Schween et al., 2003) and the kinetic models of biomass growth and recombinant protein production under the impact of phytohormone addition and bioreactor operation modes were described recently (Ruiz-Molina et al., 2022b). Further efforts have been made to select and characterize elements of gene expression regulation for the design of expression constructs encoding recombinant proteins. Various endogenous and heterologous promoter sequences were evaluated (Horstmann et al., 2004; Jost et al., 2005; Gitzinger et al., 2009), including the Physcomitrella-derived Actin5 promoter as the highest yielding promoter for molecular pharming purposes (Weise et al., 2006). In addition, secretion signals have been characterized for their effects on protein yield and secretion efficiency which facilitate downstream purification of protein products (Schaaf et al., 2005; Gitzinger et al., 2009). Finally, codon optimization of heterologous cDNAs successfully increased recombinant protein yields by preventing unwanted heterosplicing, which causes truncated target protein forms (Top et al., 2021). Another crucial element of gene expression with the potential to increase product yields are terminators. However, engineering and investigating terminators to enhance heterologous gene expression is still a neglected area compared to other yield optimization strategies.

Terminators are crucial for high recombinant protein yields by acting on transcription, pre-mRNA processing, mRNA stability and translation (Moore & Proudfoot, 2009; Zhong et al., 2023; Liu et al., 2023). At the start of transcription, the gene promoter and terminator interact in a process called gene looping. This interaction is mediated via transcription factors bound to the gene’s 5’ end and components of the polyadenylation (polyA) complex bound to the conserved polyA signal (PAS) 5’-AATAAA-3’ at the gene’s 3’ end (Tan-Wong et al., 2009; Tan-Wong et al., 2012). Gene loops facilitate RNA Polymerase II (pol II) recycling, transcriptional directionality and play a role in transcriptional memory by interacting with nuclear pores thereby facilitating transcriptional re-induction and transcript transport (Tan-Wong et al., 2009; Tan-Wong et al., 2012). The immature transcript is subsequently processed in a two-step reaction. First, the pre-mRNA is cleaved 10-30 bases downstream of the PAS (Bentley, 2005; Mandel et al., 2008). Subsequently, a polyA polymerase (PAP) adds 50 – 200 adenine residues to the polyA site, marking the end of transcription. The presence of the polyA tail is a prerequisite for the interaction with the polyA binding protein (PABP) and determines the downstream fate of the mRNA. Most transcripts undergo alternative polyadenylation (APA), a process in which a pre-mRNA is processed into multiple mRNAs that differ in their 3’UTR lengths (Guo et al., 2016; Mayr, 2016). The RNA-protein complex mediates the export into the cytoplasm and slows down mRNA degradation by 3’ exonucleases, thereby promoting mRNA stability which is linked to higher protein levels (Mandel et al., 2008). Improperly polyadenylated transcripts, in turn, are degraded faster. The polyA-bound PABP interacts with translation initiation factors present at the 5’ cap of the mRNA resulting in the formation of mRNA loops which facilitate ribosomal recycling and increase translation efficiency (Hoshino, 2012; Paek et al., 2015; Choe et al., 2018). Ultimately, ongoing de-adenylation of the polyA tail leads to PABP uncoupling and activation of the mRNA decay pathway. Hence, proper 3’end processing of transcripts is essential to ensure efficient translation.

Although members of the polyA complex and other protein factors involved in pre-mRNA 3’ processing are somewhat conserved between mammals, yeast, and plants, the RNA motifs involved differ between them (Zhao et al., 2009). In plants, polyadenylation is mediated via four main motifs, namely the cleavage element (CE) containing the cleavage site (CS) or polyA site, the near upstream element (NUE) and the far upstream element (FUE) (Rothnie, 1996; Li & Hunt, 1997; Rothnie et al., 2001; Loke et al., 2005). The CS consists of the dinucleotide motif YA (CA or UA) and is located at position −1 and −2 of the point of cleavage. The CE motif is U-rich and comprises 10 bases up- and downstream from the CS. The FUE spans 6 – 18 bases, is located 50 bases upstream of the CS and acts as an enhancer element which is less defined in its nucleotide composition but rich in U and G. The NUE locates 13 – 30 bases upstream of the CS and comprises the PAS (5’-AATAAA-3’). While highly conserved in mammals, in plants the NUE can span between 5 – 10 bases and is more diverse. In the angiosperm Arabidopsis (*Arabidopsis thaliana*) 5’-AATAAA-3’ is the major NUE but comprises only 10% of all identified motifs, whereas TGTAA is the major NUE motif in the alga *Chlamydomonas reinhardtii* (Loke et al., 2005; Wodniok et al., 2007).

Despite the important role of terminators in modulating gene expression, little attention has been paid to enhancing recombinant protein yields by analysing and engineering terminator sequences in plants. This is emphasized by the fact that the nopaline synthase (NOS) terminator from *Agrobacterium tumefaciens* and the cauliflower mosaic virus CaMV 35S terminator have been used in plant-based expression systems for decades (Pietrzak et al., 1986; Macfarlane et al., 1992). An improvement of this terminator setup was the application of the Arabidopsis heat shock protein (HSP) 90 terminator. This terminator increased the expression of a GUS reporter gene two-fold compared to the NOS terminator in transiently as well as in stably transformed plants (Nagaya et al., 2010). Terminators moved further into the spotlight resulting in a range of new insights affecting plant biotechnology. In *N. benthamiana*, an intronless version of the *N. tabacum* extensin (EU) terminator was characterized and achieved a three-fold increase in signal strength compared to its intron-containing version and a 13.6-fold increase compared to the NOS terminator (Diamos & Mason, 2018; Rosenthal et al., 2018). Combining NtEUt in chimeric double terminators with NOSt, 35St, and AtHSP90t further increased reporter levels up to 37.7-fold. Here, synergistic effects were observed between terminator pairs depending on their identity, positioning (1^st^ or 2^nd^), and heterogenicity (heterologous pairs outperformed homologous pairs). This led to the suggestion that double terminators exhibit a more stringent transcription termination signal resulting in less read-through transcripts activating RDR6-mediated silencing of the transcript (Luo & Chen, 2007; Beyene et al., 2011; Diamos & Mason, 2018). High reporter levels caused by heterologous double terminators could also be linked to enhanced interactions of the terminators with the 5’UTR and the associated machinery (Andreou et al., 2021). By characterizing 15 terminators across a range of five different promoters, Andreou et al. (2011) found synergistic interactions between promoters and terminators. However, equivalent data from non-flowering plants like mosses are lacking.

In this study, we analysed 3’UTRs from Physcomitrella and selected and characterized putative terminator candidates. We identified endogenous terminators performing equally high as the commonly employed terminator CaMV35S. We then fused high performing candidates in different combinations as double terminators and tested their performance in combination with three promoters PpActin5, CaMV35S, and NOS to further develop the biopharmaceutical production platform Physcomitrella.

## Materials & Methods

### Plant material

Physcomitrella wildtype (new species name: *Physcomitrium patens* (Hedw.) Mitt.; (Medina et al., 2019)), ecotype “Gransden 2004” (IMSC acc. No. 40001), was cultivated axenically in mineral KnopME medium 250 mg/L KH_2_PO_4_, 250 mg/L KCl, 250 mg/L MgSO_4_, 1 g/L Ca(NO_3_)_2_, 12.5 mg/L FeSO_4_, including microelements (50 μM H_3_BO_3_, 50 μM MnSO_4_ × H_2_O, 15 μM ZnSO_4_ × 7 H_2_O, 2.5 μM KJ, 0.5 μM Na_2_MoO_4_ × 2 H_2_O, 0.05 μM CuSO_4_ × 5 H_2_O, 0.05 μM CoCl_2_ × 6 H_2_O) (Reski & Abel, 1985; Egener et al.,2002; Schween et al., 2003; Decker et al., 2015). The pH was adjusted to 5.8 with KOH and media were sterilized by autoclaving. The standard growth conditions were 22 °C with a 16/8-h light/dark photoperiod and light intensity of 50–70 μmol/m^2^/s^1^. Flasks were incubated on rotating shakers with 125 rpm speed. Plant material was disrupted using an ULTRA-TURRAX (IKA, Staufen im Breisgau, Germany) at 18,000 rpm for 90 s and transferred to fresh KnopME media every seven days.

### Characterization of 3’UTRs for length, presence of introns and poly(A) sites

A fasta file containing the latest Physcomitrella genome assembly (V3.3, Lang et al., 2018) was downloaded from the Phytozome database (https://phytozome-next.jgi.doe.gov/, Goodstein et al., 2012) and the annotated UTR sequences of 75,790 transcripts were extracted based on coordinates available from the corresponding gff3 file using a custom Perl script, also available from Phytozome. Redundant UTR sequences were removed using the “delete duplicates” Excel-function. The length of each sequence was calculated using the Excel-function “LEN”. The “COUNTIF” function was used to count the number of 3’UTRs having an intron (“int”) and not having an intron (“no_int”). To predict the number of poly(A) sites in each 3’UTR the Poly(A) Site Sleuth (PASS) tool for poly(A) site prediction in plants (Ji et al., 2007; Ji et al., 2015) was used. For that purpose, all sequences smaller than 50 were removed yielding 52,952 sequences. The PASS2.0 software package for poly(A) site prediction was performed at default configurations.

### Selection of terminator candidates

The first selection of terminator candidates was based on microarray data from Hoernstein et al. (2018) (https://www.ebi.ac.uk/arrayexpress/experiments/E-MTAB-5374/files/) which included the expression levels of the transcripts of the Doko#69 line (Eleva GmbH, Freiburg, Germany, IMSC Acc. No. 40828). The translation of gene names from the V1.6 annotation to the newest V3.3 gene model (Lang, et al., 2018) was performed with the aid of the Phytozome database (Goodstein et al., 2012). The sequence of the 3’UTR for each gene, the presence of introns, the strand, and the coordinates of the 3’UTR were imported from the corresponding gff3 file, also available at Phytozome (https://phytozome-next.jgi.doe.gov/, Goodstein et al., 2012). All genes without an annotated 3’UTR were excluded from the list. For each gene containing an intron in the 3’UTR, their sequence was analysed manually on Phytozome to determine the length of the exons. Redundant UTR sequences were removed using the “delete duplicates” Excel-function. miRNA binding sites were analysed using the tool psRNAtarget (Dai & Zhao, 2011; Dai et al., 2018) containing 280 published miRNAs sequences using the pre-selected parameters (scoring scheme V2 (2017), accessed on 12^th^ of May 2020). The Gene Ontology (GO) search was performed based on GO annotation downloaded from the PEATmoss database (https://peatmoss.plantcode.cup.uni-freiburg.de/downloads/Annotations/P.patens_lookup_table_20181029.xlsx, Perroud et al. (2018), Fernandez-Pozo et al. (2020)).

### Construction of plasmids

The normalization vector 35Sp-Rluc-35St comprising the *Renilla* luciferase originates from Horstmann et al. (2004) (Supplementary file S1.1). The expression cassette consists of a long version of the CaMV 35S promoter, the *Renilla* luciferase coding sequence (CDS) and the CaMV 35S terminator. The testing vectors comprising the firefly luciferase were adapted from the 35Sp-Fluc-35St initially described by Horstmann et al. (2004) (Supplementary file S1.2). Single terminator testing vectors were obtained either by restriction-ligation cloning using restriction enzymes PstI and BglII. In that case, 35Sp-Fluc-35St was used as a template from which the 35S terminator was removed via digestion using PstI and BglII. New terminator candidates were amplified using Phusion Polymerase (Thermo Fisher Scientific Inc., Waltham, USA). PstI and BglII restriction enzyme sites were added to the template via overhang extension PCR for subsequent ligation into the firefly template vector. Alternatively, a Golden Gate-like one pot cloning reaction using type IIS enzymes Eco31I and Esp3I was used. For that purpose, the template Pro-Fluc-Ter_GG placeholder vector (Supplementary file S1.3) was obtained by removing Eco31I and Esp3I restriction sites in the template vector 35Sp-Fluc-35St. One Eco31I in the ampicillin selection marker was removed by site-directed mutagenesis via PCR. Two Esp3I restriction enzyme recognition sites were removed by deleting a 51 bp large segment in the non-coding backbone. The 35S promoter and terminator were removed and substituted by a 32 and 33 bp spanning placeholder element bearing outwards facing recognition sites for the type IIS enzymes Esp3I and Eco31I, respectively. The placeholder vector Pro-Fluc-Ter_GG and PCR amplicons for the promoter and terminator region containing Esp3I and Eco31I recognition sites, respectively, were fused with T4 Ligase, 0.5 mM ATP and the restriction enzymes in 1x Fast Digest buffer (Thermo Fisher Scientific Inc., Waltham, USA). The reaction mixture was incubated for 1 min at 37 °C and 1 min at 16 °C for in total 30 cycles. Ligated vectors were transformed into *E. coli* DH5α and positive clones were selected via Colony PCR using Taq polymerase (VWR, Radnor, USA). Correct insertion was checked via Sanger sequencing at Eurofins Genomics (Ebersberg, Germany). Primers were synthesized by Eurofins Genomics and are listed in Supplementary table S2. All vectors used in this study are listed in Supplementary table S3.

### Protoplast preparation

For protoplast preparation, existing protocols (Hohe & Reski, 2002) were adapted. Here, 100–200 mL protonema suspension from flasks cultivated in KnopME medium pH 4.5 were used as starting material (around 250 mg/L of dry weight). Plant material was disrupted as described above 7 days prior to protoplast preparation. In addition, 1 day prior to isolation of protoplasts the plant material was transferred to fresh media at pH 4.5. Moss material was incubated in the dark for 2 h in 0.5 M mannitol (adjusted to pH 5.8 with KOH, adjusted to 560 mOs using mannitol, sterilized by autoclaving) containing 1% Driselase (w/v) in the dark for 2 h. Subsequently, the moss material was passed through a 100 µm and a 50 µm sieve to separate protonema filaments from protoplasts. Recovered protoplasts were washed twice using 0.5 M mannitol solution before determining the protoplast number using a Fuchs-Rosenthal-counting chamber. Protoplasts were centrifuged and resuspended in 3 M media (0.5 M mannitol, 15 mM MgCl_2_, 0.1% MES, pH adjusted to 5.6 with KOH, osmolarity adjusted to 580 mOs using mannitol) resulting in a density of 1.2 x 10^6^ protoplasts/mL.

### Protoplast transformation

For moss protoplast transformation existing protocols (Hohe & Reski, 2002; Hohe et al., 2004; Decker et al., 2015) were adapted for usage in a 96-well format. Both firefly and *Renilla* vectors were applied in equimolar concentrations across samples within one experiment. To ensure readout values were within the linear spectrum, the ratios of firefly to *Renilla* luciferase vectors were adjusted for different experiments. For each transformation 63 µL of protoplast suspension was mixed with 25 µL of DNA in 0.1 M Ca(NO_3_)_2_ and 88 µL of 40% (w/w) PEG solution in a round-bottom 1 mL 96 deep-well plate (Nunc®, Roskilde, Denmark). The PEG solution was prepared by mixing 8 g of PEG (polyethylene glycol, MW4000) with 12 g of 3M media. The transformation mixture was incubated in a 96 well plate for 30 min on a rotary shaker at medium intensity. Protoplasts were washed twice with 500 µL 3M media, gently mixed, centrifuged for 10 min at 50 xg with slow acceleration and slow deceleration. The supernatant was discarded. Collected protoplasts were resuspended in 500 µL regeneration medium (KnopME + 50 g/L glucose and 30 g/L mannitol). Transfected protoplasts were incubated at 22 °C for 24 h in the dark and subsequently for 24h at a 16/8-h light/dark photoperiod and light intensity of 50–70 μmol/m^2^/s^1^.

### Dual Luciferase assays

Luciferase levels were detected using the Dual-Luciferase® Reporter Assay System (Promega, Madison, USA). Cells were centrifuged at 850 xg for 10 min and the supernatant was discarded. Protoplasts were resuspended in 80 µL passive lysis buffer (Promega) on a rotary shaker for 30 min at 4°C. To remove cell debris, samples were centrifuged for 1 min at 4 °C at 2250 xg. For each transformation 3x 20 µL of supernatant were transferred to a white flat-bottom 96 well-microplate (Greiner, Kremsmünster, Austria) to obtain three technical replicates. In the same sample, firefly luminescence was detected by adding 100 µL LARII reagent (Promega) and *Renilla* luciferase activity was detected by adding 100 µL Stop&Glo (Promega) reagent. Detection was performed in a POLARstar Omega microplate reader (BMG Labtech, Ortenberg, Germany) with shaking of the plate and a 1 s settling time after reagent injection and a 10 s read time. Normalized expression is reported as the ratio of luminescence from the firefly test construct to the *Renilla* calibrator. For each terminator candidate at least three independent transformations were performed and evaluated (n ≥ 3 biological replicates). For each biological replicate, the mean of three technical replicates was recorded and the resulting values were normalized to 35Sp35St.

### Statistical analysis

The firefly*-*luc/*Renilla*-luc reporter values of terminator candidates obtained were normalized to the control 35Sp35St before analysis via one-way ANOVA and Tukey HSD test using RStudio (version: 2022.07.2). Pearson correlation coefficients were calculated in Microsoft Excel 2019 between reporter values and the respective criteria such as number of polyA sites, number of polyA signals, expression values, and 3’UTR length (bp). The principal component analysis (PCA) and subsequent clustering using the package ‘Mclust’ (Fraley & Raftery, 2003) were generated using RStudio (version: 2022.07.2).

## Results

### Prevalence of alternative polyadenylation

For the Physcomitrella genome 87,533 transcripts are annotated of which 75,753 have an annotated 3’UTR (Supplementary table S4.1; Lang et al., 2018). Due to alternative splicing, several transcripts encoded from the same genomic locus can have the same 3’UTR. Here, only unique UTR sequences were kept for further analysis, resulting in 53,346 3’UTR sequences (Supplementary table S4.2). Of those unique sequences, 7,964 have at least one intron and 45,382 do not have an intron (**Fig. 1**). The average length of 3’UTRs is 654 bp and the median length is 524 bp (**Fig. 1A**). For 3’UTRs without intron the average and the median length is 542 bp and 471 bp respectively, whereas 3’UTRs with introns are more than twice as long with an average of 1,293 bp and a median of 1,114 bp (**Fig. 1B & 1C**).

**Fig. 1.**
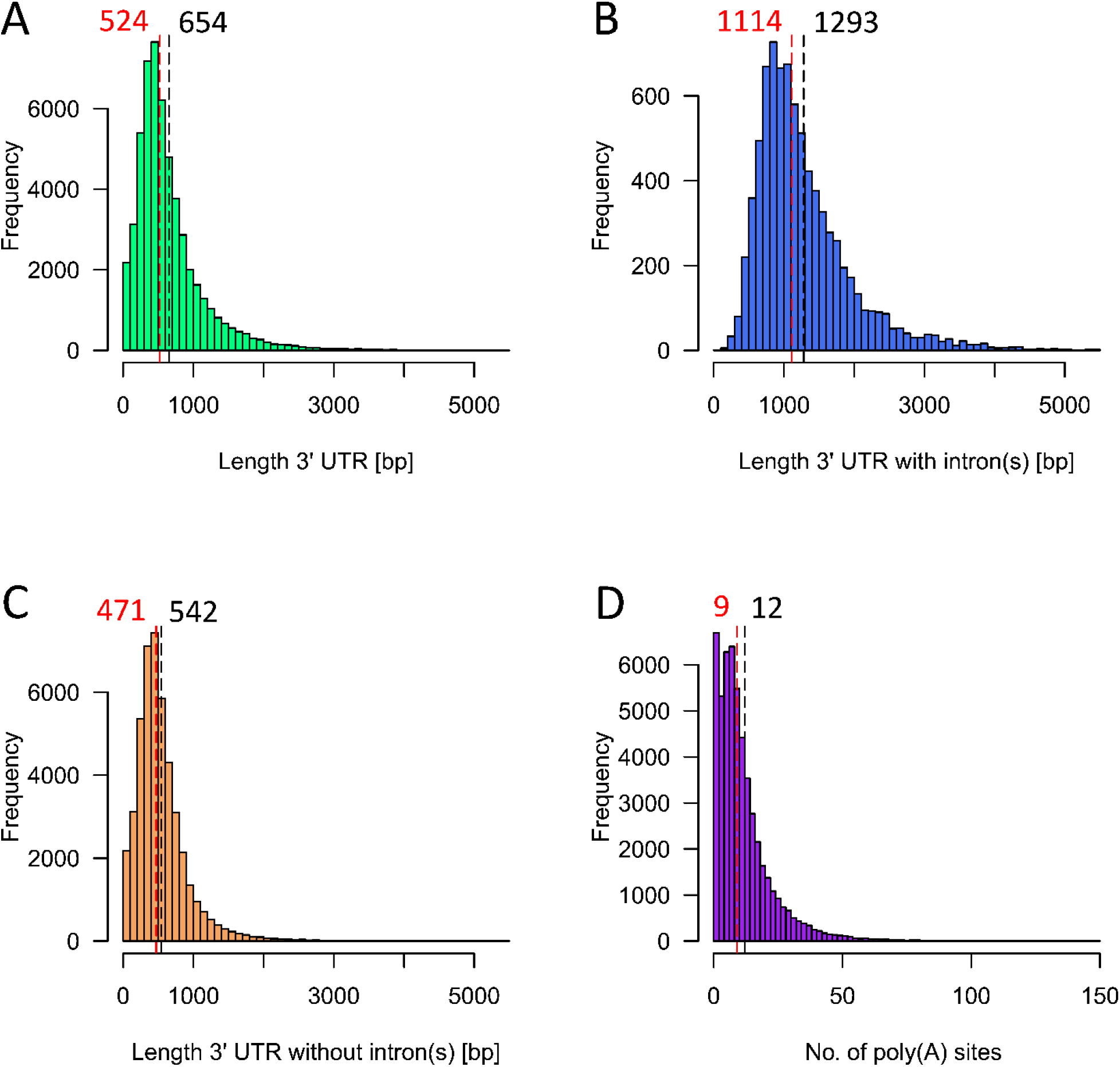
Analysis of 3’UTRs in Physcomitrella. (**A**) Length distribution of all 53,346 unique 3’UTRs. (**B**) Length distribution of unique 3’UTRs without intron(s). (**C**) Length distribution of unique 3’UTRs with intron(s). In A, B & C sequences longer than 5,500 bp were excluded. (**D**) Number of poly(A) sites predicted via PASS 2.0 software (Ji et al., 2007, 2015). For poly(A) site prediction only sequences longer than 50 bp were applicable, reducing the dataset to 52,947 sequences. The red and black dashed lines and numbers indicate the median and average, respectively.

The list was further reduced by removal of all transcripts smaller than 50 bp to allow for poly(A) site prediction. The resulting 52,947 transcripts were analysed using the PASS 2.0 software (Ji et al., 2007, 2015) to identify the location and number of poly(A) sites of each 3’UTR. Whereas around 3,607 and 1,025 transcripts exhibit no or one poly(A) site, respectively, the vast majority of 3’UTRs contain multiple poly(A) sites. In fact, >91 % of transcripts were predicted to have two or more poly(A) sites (**Fig. 1D**).

### Selection of 3’UTRs yields 14 terminator candidates

To identify endogenous terminators yielding high expression in Physcomitrella, a list of 3’UTR candidates was generated based on microarray data generated by Hoernstein et al. (2018) (Supplementary table S5.1). Here, the microarray data from the Physcomitrella Doko#69 line was utilized since this line is the platform line used by Eleva (Eleva GmbH, Freiburg, Germany) for producing biopharmaceuticals. The Doko#69 line is a double knockout for genes encoding a 1,3-fucosyltransferase and a 1,2-xylosyltransferase, which are responsible for the addition of potentially allergenic sugar residues to proteins (Koprivova, et al., 2004). Consequently, genes highly expressed in this line are considered to be highly expressed under bioproduction conditions. The 3’UTRs of the transcripts were ranked according to the highest expression average of their corresponding transcript and the top 200 were kept for further analysis. The microarray data was based on genome version 1.6 (Zimmer et al., 2013) and hence, gene IDs of the top 200 candidates were converted into version 3.3 accessions to enable further comparisons. In this step the list of candidates increased from 200 to 263, since several genes annotated in the V1.6 version had multiple matches in the V3.3 version and therefore multiple sequences (Supplementary table S5.2). After deletion of redundant sequences corresponding to splice variants within the annotation V3.3, and those without known 3’UTR sequences, 223 genes remained (Supplementary table S5.3). Long 3’UTRs are linked to mRNA decay pathway activation and shorter mRNA half-lives in Arabidopsis, *N. benthamiana*, and yeast (Kertész et al., 2006; Yamanishi et al., 2013; Srivastava et al., 2018). With respect to the determined average 3’UTR lengths shown in **Fig. 1**, a 400 bp cut off threshold was chosen yielding 65 sequences. In the case of intron-containing transcripts, the threshold was applied only to the length of the exons, resulting in one intron-containing terminator candidate (UK2, **Table 1**). Due to difficulties generated by the translation of the gene names, where some sequences of transcripts obtained for the V3.3 annotation corresponded to genes poorly expressed in the original V1.6 annotation of the microarray data, their expression in the Gransden WT line was imported from the PEATmoss expression atlas (Perroud et al., 2018; Fernandez-Pozo et al., 2020). The PEATmoss expression atlas offers expression data for tissues of different developmental stages and growth conditions. Of interest were the expression levels in protonema tissue cultivated in Knop liquid culture (Klq), the tissue and cultivation media used for bioproduction. Comparing the two expression datasets revealed that some genes highly expressed in the Doko#69 (V1.6-based microarray data), were lowly expressed in Physcomitrella WT (V3.3-based RNA-seq data, PEATmoss) (Supplementary table S5.4). Transcripts with no expression values or low values in one of those two datasets were removed from the list, resulting in 39 sequences (Supplementary table S5.5). Due to APA and other motifs potentially located further downstream of the poly(A) site (Hunt, 2008; Li & Du, 2014; Srivastava et al., 2018), 100 bp of the downstream genomic sequence were added to the selected 3’UTRs. The list of 44 3’UTRs was checked for possible miRNA binding sites employing the psRNAtarget tool (Dai et al., 2018). miRNAs are small non-translated RNA molecules which bind complementary sequences in the mRNA and consequently trigger translational inhibition or mRNA degradation (Jones-Rhoades et al., 2006; Narsai et al., 2007; Khraiwesh et al., 2010). For the set of 43 Physcomitrella miRNAs available within psRNAtarget, 52 binding sites were found on 25 different genes (Supplementary table S5.5). Among the 3’UTRs without a predicted microRNA binding site, the eight candidates with the highest expression levels in WT were selected (**Table 1**). An exception was made for a gene encoding a Chlorophyll a/b-binding protein (Cab1, Pp3c13_7900V3.1). Here, one miRNA binding site is predicted but this gene showed the highest expression in that same dataset, yielding nine candidates in total. In addition, UK2 was included as a candidate since it showed the highest expression values among intron-containing 3’UTRs. However, for molecular characterization, the first exon of 44 bp and the intron of 297 bp were removed. This decision was based on the finding that splicing events in Physcomitrella 3’UTRs are positively correlated with nonsense-mediated decay (NMD) of transcripts, a process during which aberrant transcripts are targeted for degradation (Lloyd et al., 2018). Moreover, Rosenthal et al. (2018) showed a three-fold increase in signal strength in *N. benthamiana* when removing the intron from the tobacco extensin terminator.

**Table 1.** Final selection of 18 terminator candidates for testing in Physcomitrella. Column 2 provides the sequence identifiers of Physcomitrella (no. 1 - 14) or the GenBank identifier or publication for sequences from other organisms (15 - 18). Columns 3 & 4 list gene names and the abbreviation used in this publication. Column 5 shows the expression (fragments per kilobase of transcript per million mapped reads (FPKM)) values of the candidates for the Gransden WT line cultivated in liquid Knop media (PEATmoss, Fernandez-Pozo et al., 2020). Column 6 contains information about presence of motifs, introns, miRNA binding sites predicted via psRNATarget using the list of Physcomitrella microRNAs) and poly(A) sites (predicted in PASS 2.0) (Ji et al., 2007, 2015; Dai et al., 2018). For genes Pp3c12_6700V3.1 and Pp3c16_2460V3.1 no putative gene functions are available and therefore named unknown protein and unknown protein 2.

| # | Gene Identifier/<br>Source | Gene name | Abbreviation | Expression in<br>protonema Knop<br>suspension culture<br>(FPKM) | Annotation |
| --- | --- | --- | --- | --- | --- |
| 1 | Pp3c13_7900V3.1 | chlorophyll a/b binding<br>protein | Cab1 | 5428.09 | 1x miRNA<br>binding site, 5x<br>poly(A) sites,<br>AATAAA motif |
| 2 | Pp3c13_6860V3.1 | dehydrin | dehyd | 2382.07 | 5x poly(A) sites |
| 3 | Pp3c21_4230V3.1 | plastocyanin | petE | 1178.96 | 2x poly(A) sites |
| 4 | Pp3c9_3440V3.1 | chlorophyll a/b binding<br>protein | Cab2 | 884.36 | 9x poly(A) sites |
| 5 | Pp3c22_11690V3.1 | catalase | Catalase | 879.7 | 1x poly(A) site |
| 6 | Pp3c27_2340V3.1 | photosystem II 10 kDa<br>polypeptide | PSII | 866.63 | 2x poly(A) sites |
| 7 | Pp3c12_6700V3.1 | Unknown protein | UK | 834.25 | 3x poly(A) sites |
| 8 | Pp3c3_18750V3.1 | Ribulose-bisphosphate<br>carboxylase small chain | rbcS | 749.16 | 2x poly(A) sites |
| 9 | Pp3c2_6770V3.1 | elongation factor 1-alpha | EF1A | 535.83 | 3x poly(A) sites |
| 10 | Pp3c23_22230V3.1 | photosystem I subunit IV | psaE | 529.37 | 6x poly(A) sites |
| 11 | Pp3c3_11250V3.1 | 60S ribosomal protein<br>L18A | RP-L18A | 288.67 | 5x poly(A) sites,<br>AATAAA motif |
| 12 | Pp3c11_21120V3.1 | 40S ribosomal protein<br>S6e | RP-S6e | 213.83 | 6x poly(A) sites,<br>AATAAA motif |
| 13 | Pp3c16_2090V3.1 | 40S ribosomal protein<br>S18e | RP-S18e | 213.41 | 2x poly(A) sites,<br>AATAAA motif |
| 14 | Pp3c16_2460V3.1 | Unknown protein 2 | (nol)UK2 | 82.39 | 10x poly(A)<br>sites (7 w/o<br>intron), intron,<br>4x AATAAA<br>motif (3 w/o<br>intron) |
| 15 | GenBank:<br>MN379653.1 | 35S (Cauliflower Mosaic<br>Virus) | 35S | N/A | 2x poly(A) sites,<br>1x miRNA<br>binding site,<br>AATAAA motif |
| 16 | (Rosenthal et al.,<br>2018) | extensin (*N. tabacum*) | EU | N/A | 9x poly(A) sites,<br>2x AATAAA<br>motif |
| 17 | (Andreou et al.,<br>2021) | heat shock protein 90 (*A.*<br>*thaliana*) | HSP90 | N/A | 8x poly(A) sites,<br>2x miRNA<br>binding sites,<br>3x AATAAA<br>motif |
| 18 | (Andreou et al.,<br>2021) | nopaline synthase<br>(*Agrobacterium*<br>*tumefaciens*) | NOS | N/A | 3x poly(A) sites,<br>AATAAA motif |

In parallel, a second approach to identify terminator candidates was based on gene ontology (GO). Yamanishi et al. (2013) performed a genome-wide activity assessment in the yeast *Saccharomyces cerevisiae* and found terminators from genes coding for ribosomal proteins yielded high expression of transgenes. Accordingly, a search using the GO terms “ribosome”, “structural constituent of ribosome”, and “ribosome biogenesis” yielded 2,109 matches in Physcomitrella (Supplementary table S6.1). Many genes matched two or three of the categories. Duplicates were removed, resulting in a list of 1,107 unique genes (Supplementary table S6.2) of which only 3’UTRs with a length between 150 bp and 401 bp were kept, resulting in 358 genes (Supplementary table S6.3 and S6.4). Subsequently, the presence of a 5’-AATAAA-3’ signal was checked resulting in 47 3’UTRs (Supplementary table S6.5). Since the PAS is an important element for mRNA stability and half-life (Rothnie, 1996), this step was meant to isolate genes with a higher chance of being correctly processed and subsequently more stable. After excluding 3’UTR transcripts which were redundant or not expressed in the Gransden WT protonema liquid culture, imported from PEATmoss, 19 genes remained (Supplementary table S6.6). Using the psRNAtarget tool (Dai et al., 2018), the six 3’UTRs with the highest expression were checked for miRNA binding sites (Supplementary table S6.7). Finally, three 3’UTRs without miRNA binding sites were selected as terminator candidates (**Table 1**).

Lastly, the 3’UTR of a Chlorophyll a/b-binding protein (Cab2, Pp3c9_3440V3.1) was added to the list. This UTR is 511 bp long and therefore exceeds the length threshold of 400 bp introduced earlier. However, Cab2 does not contain predicted miRNA binding sites, exhibits the 4^th^ highest expression values compared to other selected terminator candidates (**Table 1**), and contains nine predicted poly(A) sites. To enable comparison of the endogenous terminator candidates the widely used benchmark terminators Ca35St, NtEUt, AtHSP90t, and NOSt were included resulting in a final list of 18 different terminators (**Table 1**).

### Endogenous terminators yield as equally high as the CaMV 35S terminator

To systematically evaluate the 18 individual terminator candidates selected (**Table 1**) we performed Dual Luciferase assays. Consequently, we cloned expression constructs containing the firefly luciferase CDS under control of the 35S promoter and the respective terminator candidates. Activity of the transiently expressed reporters was measured 48 hours after PEG-mediated transfection of these vectors in Physcomitrella protoplasts. To allow for reliable screening of large vector libraries, a 96 well plate-based transformation protocol for Physcomitrella protoplasts was established. Differences in transformation efficiency between samples were normalized with the activity of the co-transfected *Renilla* luciferase under control of the 35S promoter and 35S terminator.

From the tested single terminators, eight candidates showed no difference to the reference terminator 35S, including the heterologous terminators AtHSP90, NOS, and NtEU (**Fig. 2**). In addition, Physcomitrella-derived terminators PpRPS6et, PpnoIUK2t, PpCab2t, PpCab1t, and PppetEt performed as well as 35St. However, several endogenous terminators showed a significantly lower performance compared to 35St. Furthermore, there was no significant difference in the performance of the NOS terminator and the NtEU terminator (**Fig. 2**, NOSt, NtEUt). This is in contrast to Rosenthal et al. (2018), who found NtEUt to increase reporter levels by 13.3-fold in *N. benthamiana* as compared to NOSt (Diamos & Mason, 2018; Rosenthal et al., 2018). The lowest performing terminator is PpUK.7t at 0.14, 7.9 times lower than the highest performing terminator AtHSP90t. We could identify Physcomitrella endogenous terminators performing as well as the widely used 35S terminator. Subsequently, we tested combinations of our best performing terminators in double terminator constructs.

**Fig. 2.**
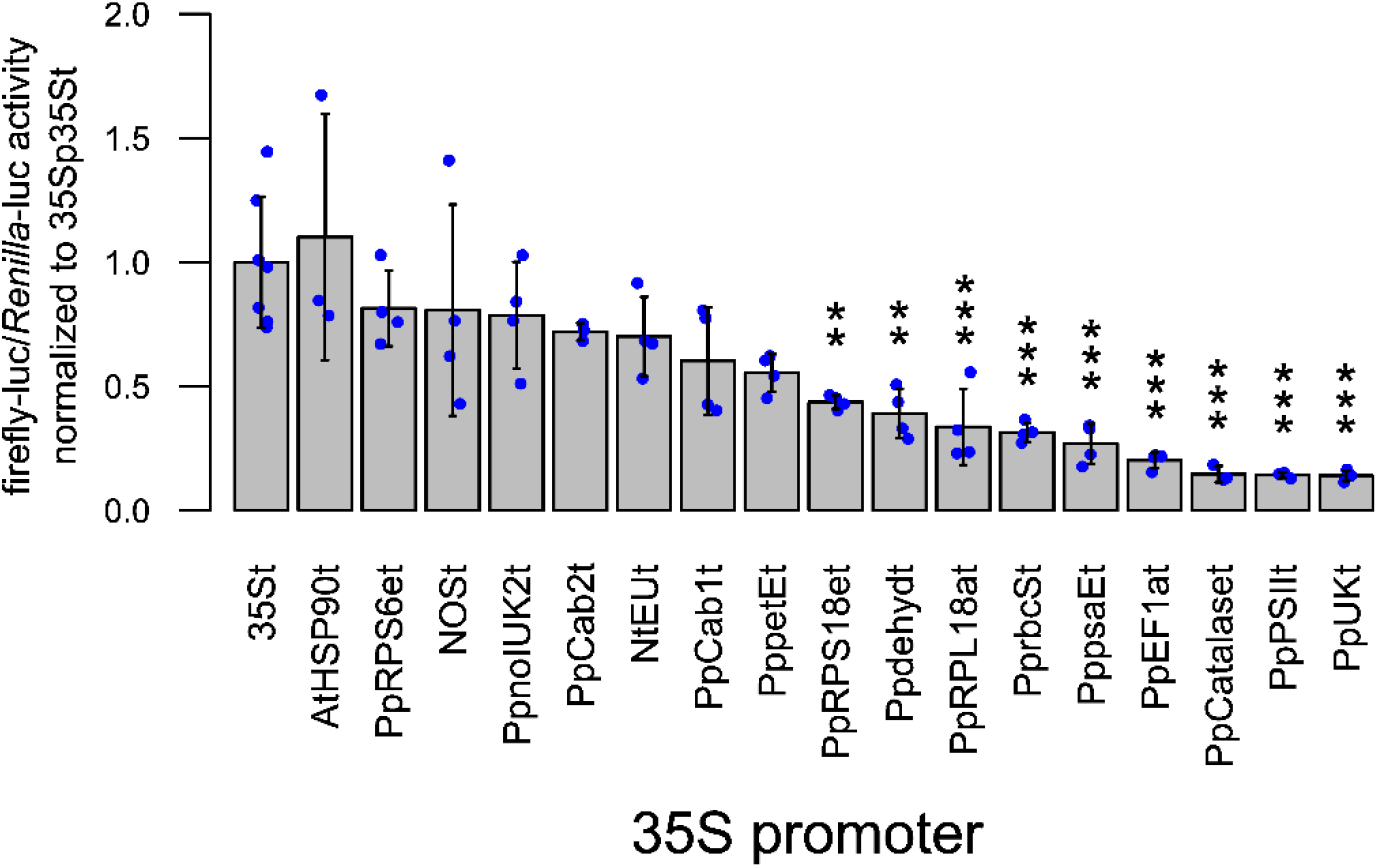
Characterization of single terminator candidates. Terminator candidates (Table 1) were cloned into a vector carrying the firefly luciferase coding sequence and the 35S promoter. Terminators were characterized via Dual Luciferase assay and obtained firefly-luc/*Renilla*-luc reporter values of single candidates normalized to the 35S terminator. Blue dots represent means of three technical replicates of a biological replicate. For each terminator n≥ 3 biological replicates were analysed. Bars represent means of all biological replicates with standard deviations. Significance levels are based on a one-way ANOVA (*p*=7.91E-11, α=0.05) with subsequent Tukey HSD post-hoc test. Indicated differences are between 35St and the respective terminator (*p<0.05; **p<0.005; ***p<0.0005).

### Endogenous double terminators up to three-fold more efficient than respective single terminators

Combining two terminators in tandem in one construct was shown to increase reporter signal levels in various angiosperms, including Arabidopsis and *N. benthamiana* (Luo & Chen, 2007; Beyene et al., 2011; Yamamoto et al., 2018; Diamos & Mason, 2018). These synergistic effects were observed between terminator pairs depending on their identity, positioning (1^st^ or 2^nd^), and heterology (heterologous pairs outperformed homologous pairs). This raised the question if different combinations of double terminators would outperform the respective single terminators in Physcomitrella. To address this, we generated different combinations of double terminators and tested them using the same experimental setup employed to test the single terminators. First, we generated homologous and heterologous combinations of the two benchmark terminators 35St and NOSt. Likewise, we generated combinations of the endogenous terminators PpCab2t and PppetEt. Finally, we generated combinations of a heterologous and an endogenous terminator, AtHSP90t and PpRPS6et. The combinations of 35St and NOSt did not show any improvement compared to the respective single terminators, independent of how they were combined (**Fig. 3A**). In contrast, Diamos & Mason (2018) observed an 18.4-fold increase in GFP reporter signal levels between 35St-NOSt double terminator and NOSt alone in *N. benthamiana*. Beyene et al. (2011) observed an up to 65-fold increase in EYFP levels between 35St-NOSt and 35St in sugarcane (*Saccharum officinarum*). In contrast, all PpCab2t and PppetEt double terminator combinations outperformed the single terminator petEt in Physcomitrella (**Fig. 3B**). The combination PpCab2t-PppetEt even outperformed both single terminators, yielding around 2.3-fold and 2.9-fold higher than PppetEt and PpCab2t, respectively. PpCab2t-PppetEt also performed higher than the homologous double terminators and even the reciprocal combination PppetEt-PpCab2t. These observations are in line with the findings of Diamos & Mason (2018) regarding the role of identity and positioning of tandem terminators, although the fold changes observed here with an enzymatic reporter were again lower than the fold changes observed in terminator combinations in *N. benthamiana* with a fluorescent reporter.

**Fig. 3.**
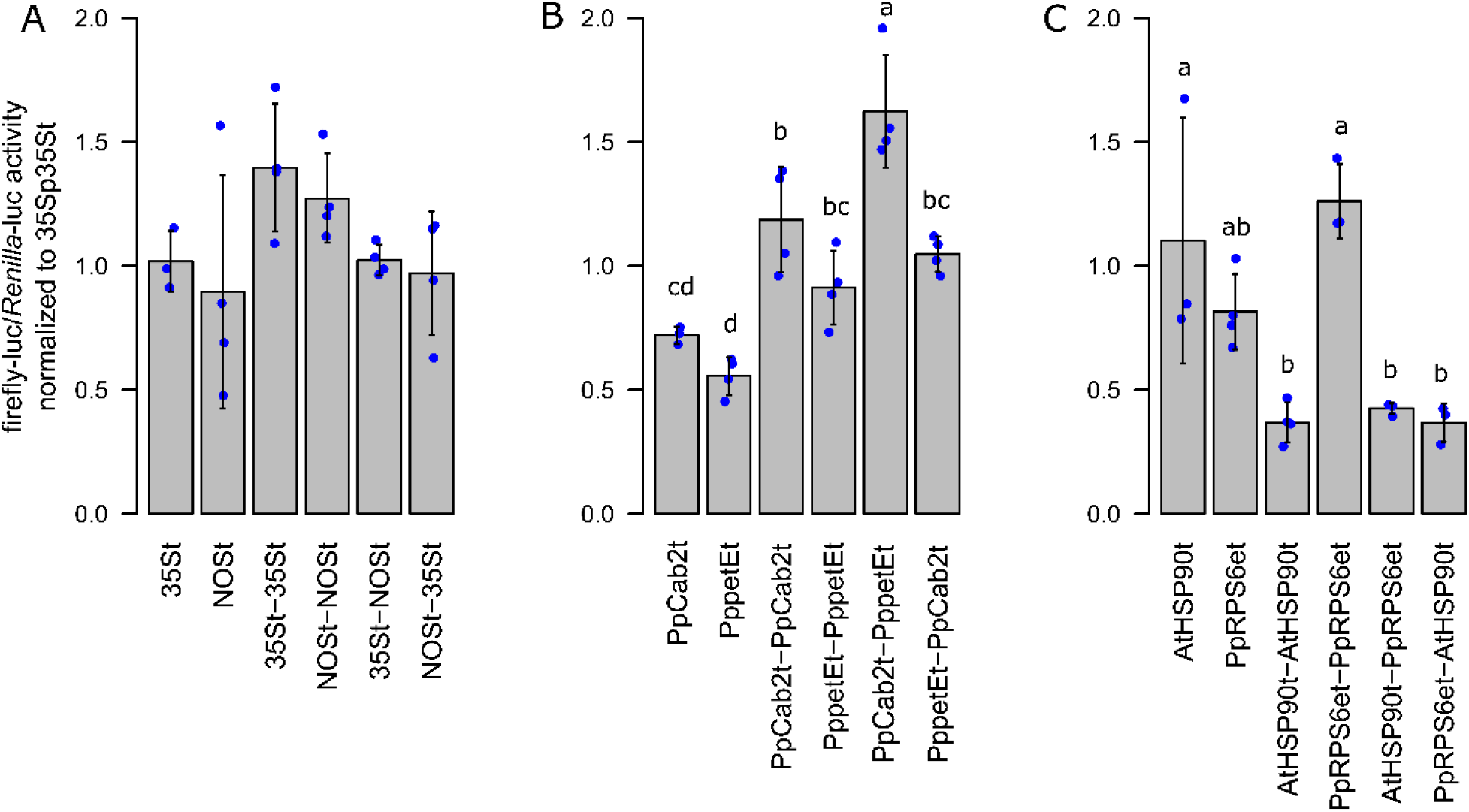
Characterization of double terminators. Single terminators were fused in homologous and heterologous pairs in vectors bearing the firefly luciferase coding sequence and the 35S promoter. Double terminators and their respective single terminators were characterized via Dual Luciferase assays and obtained firefly-luc/*Renilla*-luc reporter values normalized to the 35S terminator. Blue dots represent means of three technical replicates of a biological replicate. Bars represent means of all biological replicates (n≥3) with standard deviations. No significant difference in performance were observed for combinations of the 35St and NOSt (**A**, One-way ANOVA, p=0.11, α=0.05). Significant differences in performance were observed in combinations of PpCab2t and PppetEt (**B**, One-way ANOVA, p=4.16E-7, α=0.05) and in combinations of AtHSP90t and PpRPS6et (**C**, One-way ANOVA, p=8.42E-05, α=0.05). Significance levels and compact letter display are based on a one-way ANOVA with subsequent Tukey HSD post-hoc test.

Moreover, terminator combinations based on AtHSP90t and PpRPS6et depict another deviation from findings in other plant systems. Here, combinations either did not have any deviating effect on performance compared to the respective single terminators, or it decreased performance significantly (**Fig. 3C**). All terminator combinations containing AtHSP90t yield lower than the respective single terminator, indicating negative interactions between AtHSP90t and PpRPS6et or itself in Physcomitrella. Such a negative effect of AtHSP90t in double terminators was not observed by Yamamoto et al. (2018) when testing AtHSP90t in combination with NtEUt and 35St in *N. benthamiana*.

Taken together, double terminators can have a positive effect on gene expression in Physcomitrella. These effects depend strongly on the identity and positioning of the pairs. However, no clear pattern can be deduced at the moment. Also, the performance of certain double terminator combinations, e.g. 35St-NOSt, seems to vary from species to species.

### Promoters are more dominant in increasing gene expression levels than terminators

In Physcomitrella, several promoters were characterized (Horstmann et al., 2004; Jost et al., 2005; Weise et al., 2006; Gitzinger et al., 2009) and PpActin5p, 35Sp and NOSp are among the most employed promoters (Top et al., 2019; Bohlender et al., 2020; Ruiz-Molina et al., 2022a). Data generated in Arabidopsis indicates that a terminator’s performance is dependent on the promoter it is paired with, suggesting that performance depends more on combinatorial than on additive effects (Andreou et al., 2021). Consequently, the performance of terminators would not be constitutive and must be regarded in relation to the promoter employed. In this respect, we further analysed the performance of selected terminators in combination with the PpActin5 and the NOS promoter, respectively. Again, reporter values were obtained by normalizing to 35Sp35St which was set to the value 1.

Across the three promoters, no striking difference in terminator performance depending on the selected promoter was observed (**Fig. 2, 4A, 4B**). However, our data reveal the dominant role of the promoter in determining gene expression levels in Physcomitrella. As already seen for the 35S promoter (**Fig. 2**), all tested terminators, except PpnoIUK2t paired with PpActin5p, performed as well as the 35S terminator (**Fig. 4A, 4B**). For the NOS promoter the terminators 35St, Cab2t, dehydt, and AtHSP90t perform significantly better than noIUK2t. In the case of the PpActin5 promoter no significantly different performing pair could be identified. This can be explained by the generally large standard deviation across all terminators combined with the PpActin5 promoter (**Fig. 4B**). This standard deviation is lower for the 35S promoter (**Fig. 2**) and even lower for the NOS promoter (**Fig. 4A**). However, terminators NOSt and AtHSP90t exhibit an enlarged variation across all three promoters. The only terminator suggesting the presence of promoter-terminator interactions is noIUK2t. Terminator noIUK2t yields significantly higher than rbcSt when combined with the 35S promoter (**Fig. 2**) but shows no difference to rbcSt when combined with the NOS or PpActin5 promoter (**Fig. 4A, 4B**). Still, apart from this observation, our data does not indicate promoter-terminator interactions to be a prevalent phenomenon in Physcomitrella.

**Fig. 4.**
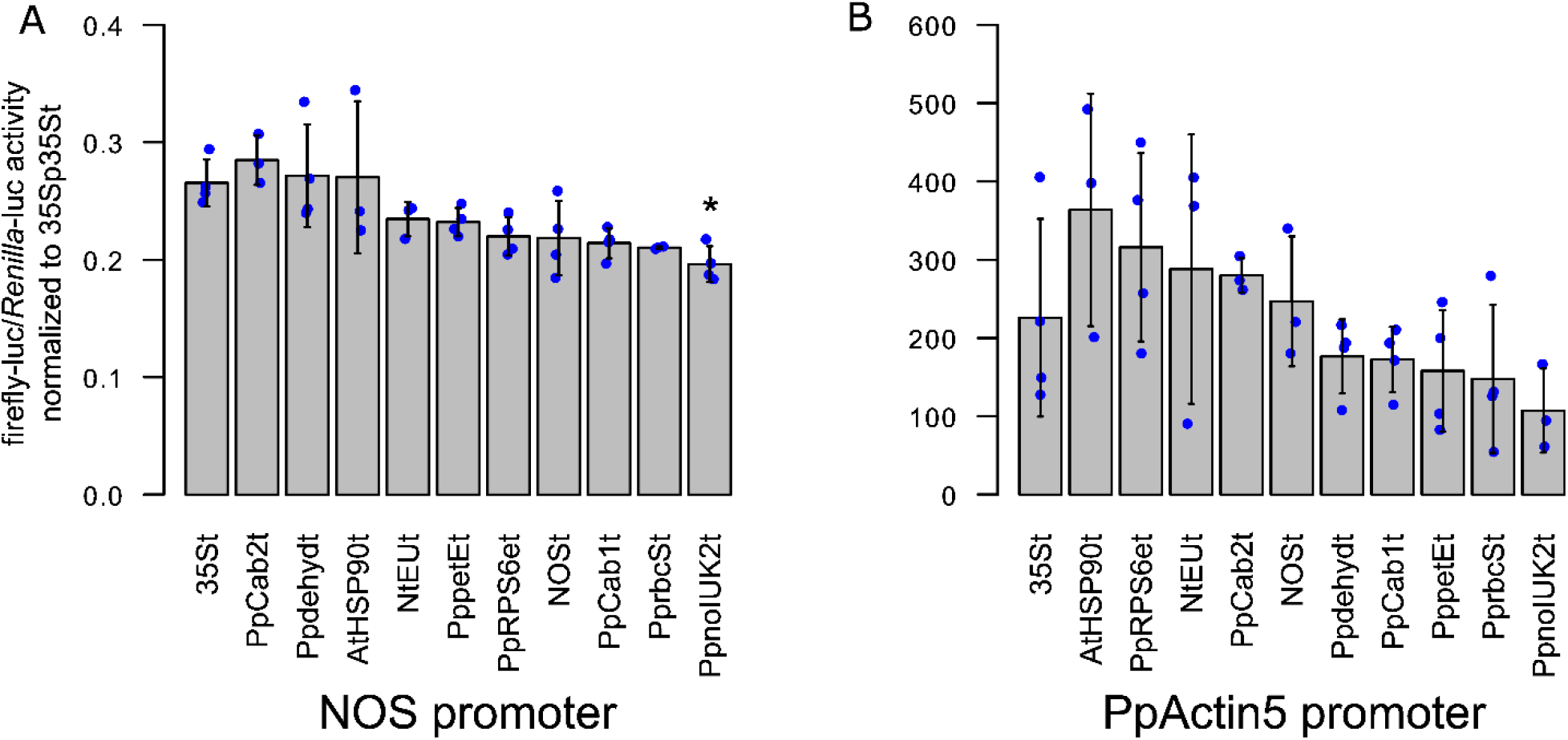
Characterization of terminators in combination with the NOS or the PpActin5 promoter. A selection of single terminators were cloned in vectors bearing the firefly luciferase coding sequence and either the (**A**) NOS promoter or the (**B**) PpActin5 promoter. The promoter-terminator combinations were characterized via Dual Luciferase assays and obtained firefly-luc/*Renilla*-luc reporter values normalized to a testing vector bearing the 35S promoter and 35S terminator. Blue dots represent means of three technical replicates of a biological replicate. Bars represent means of all biological replicates (n≥3) with standard deviations. In combination with the NOS promoter, only terminator PpnoIUK2t shows significant difference in comparison to 35St (A, One-way ANOVA, p=0.0012, α=0.05). For the PpActin5 promoter no tested terminator significantly differed in performance from 35St (B, One-way ANOVA, p=0.046, α=0.05). Significance levels are based on a one-way ANOVA with subsequent Tukey HSD post-hoc test. Indicated differences are between 35St and the respective terminator (*p<0.05).

Instead, our data clearly shows a dominant role of the promoter compared to the terminator when aiming at optimizing gene expression levels. The fold change from lowest to highest performing terminator when using the 35S promoter is 7.9 when comparing 18 different terminator candidates (**Fig. 2**). For the NOS and PpActin5 promoters the fold changes are 1.5 and 3.4, respectively, when comparing 11 different terminator candidates (**Fig. 4A, 4B**). In turn, AtHSP90t yields 0.27 if combined with the NOS promoter, 1.1 with the 35S promoter, and 363.63 with the PpActin5 promoter, an increase of about 1,350-fold. These results clearly underline the importance of choice of promoter over terminator when optimizing gene expression levels. Further, it confirms the superiority of the PpActin5 promoter as compared to the 35S or NOS promoter for recombinant protein production in Physcomitrella (Weise et al., 2006).

### Performance is determined by poly(A) signals and sites

Finally, we investigated which of our terminator selection criteria influenced terminator reporter levels the most. For that purpose, Pearson correlation coefficients were calculated between reporter values and different terminator attributes (**Table 2**). These attributes represent criteria that were applied during terminator selection, e.g., the number of 5’-AATAAA-3’ PAS motifs, expression levels from the PEATmoss database (Fernandez-Pozo et al., 2020), and 3’UTR length. In addition, the number of poly(A) sites as predicted via the PASS2.0 software (Ji et al., 2007, 2015) was included, even though this criterion was not applied during terminator candidate selection. Expression values originate from the Physcomitrella Gransden wildtype line as seen in **Table 1**. Since no expression values are available for the non-Physcomitrella derived terminators 35St, NOSt, AtHSP90t, and NtEUt; these heterologous terminators were excluded from correlation calculations between reporter levels and expression values.

**Table 2.** Pearson correlation coefficients between reporter values and terminator attributes. Pearson correlation coefficients were calculated between the reporter values obtained and the terminator attributes such as the number of PAS, number of poly(A) sites, expression levels and terminator length. The analysis was performed for reporter values obtained with each promoter. For the terminators AtHSP90t, 35St, NtEUt and NOSt expression values were not available.

|  | # PAS<br>5'-AATAAA-3' | # poly(A) sites | expression levels | length (bp) |
| --- | --- | --- | --- | --- |
| **35Sp** | 0.71 | 0.50 | 0.11 | 0.05 |
| **Act5p** | 0.27 | 0.51 | -0.17 | 0.33 |
| **NOSp** | -0.20 | 0.27 | 0.04 | 0.35 |

We found a correlation of r = 0.71 between the number of PAS and terminator performance regarding the 35S promoter. In turn, there is only a weak correlation between PAS and reporter values obtained via the PpActin5 promoter (r = 0.27) and a weak negative correlation for the NOS promoter (r = −0.20). The number of PAS was used as a terminator selection criterion only during the GO term-based approach through which candidates RP-L18a, RP-S6e, and RP-S18e were identified. The presence of PAS did not influence the selection of any other terminator candidates. Still, all non-Physcomitrella derived terminators as well as noIUK2t and Cab2t exhibit the 5’-AATAAA-3’ motif nontheless (**Table 1**). Regarding the number of poly(A) sites, terminators characterized via the 35S and PpActin5 promoter show a moderate correlation of r = 0.5 and r = 0.51 whereas for the NOS promoter only a weak correlation of r = 0.27 was observed. For expression levels no or only weak correlation was found across reporter values obtained with either of the three promoters. This finding is remarkable since expression levels were a major criterion in terminator candidate selection. Terminator candidate length exhibits a weak correlation in the PpActin5 and NOS promoter datasets, but no correlation if terminators were combined with the 35S promoter.

Since no or only moderate correlation between the selected single features was observed, we performed principal component analysis (PCA, **Fig. 5**) to investigate whether combinatorial effects on gene expression can be observed. Expression levels were excluded from this analysis for two reasons: First, the initial Pearson correlation showed no correlation between expression levels and reporter values for either the 35S, NOS or Actin5 promoter (**Table 2**). Second, expression levels were obtained from the PEATmoss database (Perroud et al., 2018; Fernandez-Pozo et al., 2020) and are only available for the transcripts of Physcomitrella endogenous terminators but not for 35St, NOSt, AtHSP90t, and NtEUt. Consequently, these terminators would be excluded from the analysis, thereby minimizing the dataset and thus the validity of the outcome. For the PCA, reporter values for each promoter series were normalized to the promoter series own combination with the 35S terminator, e.g. all reporter values obtained for terminators in combination with the Actin5 promoter were normalized to the sample Act5p35St (Supplementary table S1.2) to avoid overrating of the strong promotor effects.

**Fig. 5.**
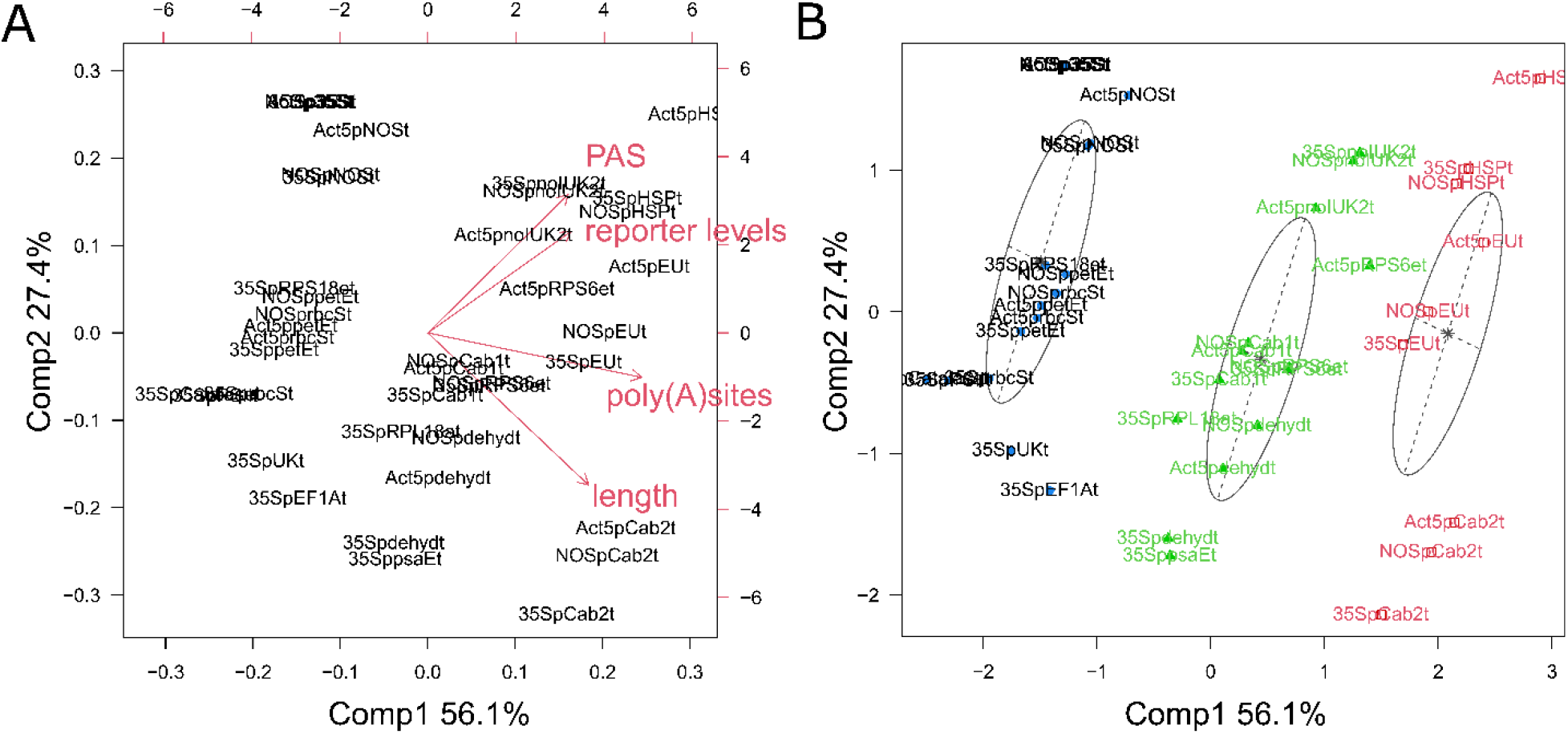
Principal Component Analysis of reporter values and terminator attributes. A PCA was calculated for the reporter values obtained, the number of PAS, the number of poly(A) sites, and terminator lengths. Reporter values of the single promoter series (PpActin5p, 35Sp, NOSp) were normalized to the respective promoter combined with the 35S terminator to remove promoter effects from the dataset. The biplot in **A** shows the different loading vectors of reporter values and terminator attributes. The PCA plot in **B** shows the result of the model-based clustering using Mclust (Fraley & Raferty, 2003). Reporter values correlate with the number of PAS whereas the number of poly(A) sites and terminator length determine cluster formation. Promoter-terminator combinations are sorted into three clusters in black, green, and red.

The biplot depicts the single data points of promoter-terminator combinations and how these scores are distributed over components (PC) 1 and 2 (**Fig. 5A**). In addition, the different terminator attributes reporter levels, the number of PAS, the number of poly(A) sites, and length are represented as loading vectors. The length of a vector represents how strongly it impacts the underlying PC whereas the angle between vectors provides information about the degree of correlation, e.g. a small angle signifies correlation whereas an angle of 90° signifies a lack of correlation. Using the PCA scores obtained for each promoter-terminator combination we performed model-based clustering using Mclust (Fraley & Raftery, 2003). Cluster uncertainties for each datapoint are available in Supplementary table S7.

The biplot in **Fig. 5A** shows that PC 1 and 2 explain 56.1% and 27.4% of variation, respectively, totalling 83.5%. The number of poly(A) sites has the strongest impact on PC 1 whereas the attributes reporter levels, number of PAS, and length are influencing PC 1 and PC 2 at varying levels. According to this PCA, the length of the terminator and the number of poly(A) sites have the strongest impact on the variance of the data but do not correlate with the obtained reporter values (**Fig. 5A**). In contrast, the number of PAS correlates with the obtained reporter values.

The PCA plot in **Fig. 5B** yields three clusters (Supplementary table S7). The clusters are orientated alongside the loading vectors reporter levels and number of PAS and orthogonal to the loading vectors length and number of poly(A) sites. Accordingly, length and number of poly(A) sites are the main attributes by which clusters were formed. In the red cluster, terminators possess 8 – 9 poly(A) sites and are on average 518 bp long. Terminators in the green cluster possess 5 – 7 poly(A) sites and have an average length of 458 bp. In the black cluster there are terminators with 1 – 3 poly(A) sites and 304 bp in length. In conclusion, none of the attributes alone is a strong determinant for terminator performance in Physcomitrella. Instead, different factors act to varying degrees on obtained reporter values. Mainly, the number of PAS but also to some degree the number of poly(A) sites influences terminator performance whereas the terminator length shows a minor effect.

## Discussion

Over decades the field of molecular pharming and plant biotechnology in general relied on a small range of terminators, namely 35St, NOSt, and AtHSP90t. However, in recent years, the selection and characterization of new terminators as a means to further increase product yields moved into the spotlight. For *N. benthamiana*, NtEUt was characterized and shown to have a wide applicability across different plant species (Rosenthal et al., 2018). Further, NtEUt and other terminators were shown to yield higher expression levels when combined as double terminators (Diamos & Mason, 2018). Terminator identity and positioning in these pairs provided insights into the mode of action behind double terminators with links to promoter-terminator interactions (Andreou et al., 2021). In this study, 3’UTRs of the production host Physcomitrella were analysed and putative terminator candidates selected and characterized. We further selected high performing candidates for double terminators and tested different combinations and positioning for their effects on gene expression. To address the hypothesis of promoter-terminator interactions we combined a range of terminators with different promoters.

The 3’UTRs of the moss Physcomitrella were analysed and found to be on average 654 bp long with most sequences at around 400 bp in length. In comparison, 3’UTRs have an average length of 242 bp in Arabidopsis and 469 bp in rice (Srivastava et al., 2018). The 3’UTRs of the alga *C. reinhardtii*, in turn, have an average length of 595 bp (Shen et al., 2008a), considerably longer than in Arabidopsis and rice and more in line with our findings from Physcomitrella. Further, we found more than 48,000 Physcomitrella 3’UTRs to exhibit one or more poly(A) sites whereas only in 3,607 3’UTRs no poly(A) site could be identified. While this finding is in line with data from other plants demonstrating the prevalence of APA in plants, Physcomitrella might be more prone to APA than other plants. In Arabidopsis, more than 70% of genes contain >1 poly(A) site according to sequencing the junction of 3’UTRs and poly(A) tails (Wu et al., 2011). In rice about 50% of genes have >1 poly(A) sites and 13% have ≥4 poly(A) sites (Shen et al., 2008b). In *C. reinhardtii* the number of genes influenced by APA decreases even further to about 33% (Shen et al., 2008a).

We used five determinants to select Physcomitrella endogenous terminator candidates from a list of 3’UTRs transcripts. These determinants were i) transcript abundance based on microarray and RNAseq data, ii) transcript length, iii) absence of miRNA binding sites, iv) presence of the PAS 5’-AATAAA-3’, and v) a GO term search using the term ‘ribosome’. The abundance of a transcript (i) derives from the balance in the interplay of transcription rate and turnover rate. A main function of terminators is to stabilize mRNAs by facilitating proper polyadenylation which prevents degradation through 3’ exonucleases and prolongs the activation of the mRNA decay pathway. Therefore, transcript abundance was considered an interesting characteristic to decipher the effect of terminators on Physcomitrella mRNA stability. Other determinants (ii - iv) were chosen which influence mRNA stability according to studies performed in Physcomitrella, plants in general, and other systems. In *N. benthamiana*, long 3’UTRs and 3’UTR-located introns were found to activate NMD of mRNAs (Kertész et al., 2006). In Arabidopsis, NMD can act as a viral restriction mechanism by targeting long 3’UTRs variants of polycistronic ORFs of potato virus X (Garcia et al., 2014). These findings are in line with data obtained in yeast (*S. cerevisiae*) showing least active terminators to be linked to longer 3’UTRs. In contrast, a genome-wide analysis of Arabidopsis NMD rates and their determinants showed 3’UTRs longer than 300 bp and the presence of introns in the 3’UTRs to not influence mRNA half-lives (Narsai et al., 2007). In Physcomitrella, not 3’UTR length but splicing within the 3’UTR, especially if close to the stop codon, is a positive predictor of NMD (Lloyd et al., 2018). As a result, we chose a relatively high 3’UTR cut-off value of 400 bp due to the ambiguous findings regarding the effect of 3’UTR length on mRNA as well as the higher average length of Physcomitrella 3’UTRs compared to other plants. Even though not actively selected against, the cut-off at 400 bp removed all but one intron-containing terminator candidate from the list (UK2t) (**Table 1**). Still, for characterization of UK2 the intron-less version (noIUK2t) was used, based on the findings of Lloyd et al. (2018), showing that splicing in the 3’UTR correlates with NMD in Physcomitrella. In addition, the intron-less NtEUt yielded higher reporter levels compared to the intron-containing version in *N. benthamiana* (Rosenthal et al., 2018). Further, an over-representation (CAGTTGAAATTT and GTGAAAVTTTTC) and under-representation (CCAACATCAT, CTTGGCTA, and THTCAWGGGT) of certain motifs in NMD-targeted transcripts was found (Lloyd et al., 2018). In Arabidopsis, destabilizing motifs were generally rich in T whereas stabilizing motifs were rich in A, including the conserved PAS motif 5’-AATAAA-3’ (Loke et al., 2005; Narsai et al., 2007) used for terminator selection in this study. For both Physcomitrella and Arabidopsis the authors state that in many cases transcripts contained motifs of both kinds, suggesting a yet unclear interplay of motifs and their effect on transcript stability. Further, the sequence identity of the most prevalent PAS in Physcomitrella has not been identified and could vary substantially from the Arabidopsis PAS as is the case for *C. reinhardtii* (PAS: TGTAA) (Wodniok et al., 2007). Moreover, studies across yeast, humans, and plants demonstrated mRNA half-lives to be linked to the biological process of the encoded protein (Wang et al., 2002; Yang et al., 2003; Narsai et al., 2007; Yamanishi et al., 2013). mRNAs encoding for proteins involved in central metabolism, energy, protein synthesis, and subcellular localization showed significantly higher half-lives in Arabidopsis as compared to transcripts classified in highly regulated processes such as stress response-related transcription factors and kinases (Narsai et al., 2007). Accordingly, many of the eleven Physcomitrella terminator candidates not selected via GO term enrichment analysis are also linked to plant central metabolism and photosynthesis (**Table 1**). miRNAs were found to negatively impact mRNA half-lives due to their established role in complementary RNA targeting, cleavage, and degradation (Jones-Rhoades et al., 2006; Narsai et al., 2007) and were therefore used as a determinant for terminator candidate selection.

Using these parameters for 3’UTR selection, a terminator candidate list was generated containing 14 Physcomitrella-derived terminators (**Table 1**). That list was further extended by adding the four heterologous terminators 35St, NtEUt, AtHSP90t, and NOSt. All terminators were characterized in combination with the 35S promoter using the Dual Luciferase system and transient protoplast transformation. Heterologous terminators NtEUt, AtHSP90t, and NOSt, as well as endogenous terminators RPS6et, noIUK2t, Cab2t, Cab1t, and petEt, performed as well as the default terminator 35St. All other tested candidates yielded less while no terminator performed significantly better than the 35St. As NtEUt, NOSt, and AtHSP90t are among the highest yielding terminators in Physcomitrella, this demonstrates the versatility of some gene expression elements across plant lineages (moss, angiosperm). Across three different angiosperms, namely *N. benthamiana*, *N. tabacum*, and lettuce, a slightly varying but nevertheless strong effect of NtEUt on reporter expression levels could be demonstrated (Diamos & Mason, 2018). Further, the functionality of NtEUt and NtEUt-based double terminators in a range of angiosperm species and tissues, such as lettuce (*Lactuca sativa*), eggplant (*Solanum melongena*), tomato (*Solanum lycopersicum*) fruits and leaves, hot peppers (*Capsicum frutescens*), melons (*Cucumis melo*), orchids (*Phalaenopsis Aphrodite*), and roses (*Rosa* sp. ‘Bonheur’) could be shown (Yamamoto et al., 2018). However, reporter levels were not quantified in that study. Here, we demonstrated EUt from *N. tabacum* to be functional in Physcomitrella. However, an increase in gene expression for NtEUt as compared to 35St and NOSt in *N. benthamiana* (Diamos & Mason, 2018; Rosenthal et al., 2018) could not be obtained in Physcomitrella. This may hint at differences between mosses and angiosperms in the use of terminators for the regulation of gene expression.

A selection of terminators was further characterized using the promoters NOSp and PpActin5p (**Fig. 4**). Here, relatively small fold changes from lowest to highest performing terminator of 1.5 and 3.4 for NOSp and PpActin5p, respectively, were found. As for the 35S promoter, heterologous terminators NtEUt, AtHSP90t, and NOSt, as well as the majority of Physcomitrella endogenous terminators performed as well as the default terminator 35St. However, large fold changes in reporter levels between the promoters NOSp, 35Sp, and PpActin5p were obvious. We further found the effect of the promoter to be independent of the paired terminator. In Arabidopsis, synergistic effects between promoters and terminators were reported by characterizing 15 terminators in combination with five different promoters (Andreou et al., 2021). They found AtHSP90t to perform consistently high, AtrbcS2b to perform consistently low, and NOSt to perform in dependence of the paired promoter. Here, we tested at least eleven terminators across three promoters and could not identify clear promoter-terminator interactions in Physcomitrella. Indeed, terminators such as NOSt and noIUK2t show varying performance across promoters in this moss. Further, AtHSP90t and PprbcSt also performed consistently high or low, respectively, across promoters. This finding is similar to data described by Andreou et al. (2021) for AtHSP90t and AtrbcS2b. Overall, the data obtained here indicate that in Physcomitrella the choice of terminator does have a huge impact on gene expression levels. Instead, the selection of the promoter has a larger impact and dominates the influence in gene expression in our assay.

After characterizing the selected terminator candidates via the Dual Luciferase system, we aimed to clarify how the selection criteria influenced terminator performance the most. For that purpose, we calculated Pearson correlation coefficients between reporter values of each promoter series and the number of PAS, the number of poly(A) sites, expression levels from the PEATmoss database (Perroud et al., 2018; Fernandez-Pozo et al., 2020) and terminator length (**Table 2**). Likewise, a PCA was performed for which expression levels were excluded (**Fig. 5**). The results indicate that the number of PAS and the number of poly(A) sites influence terminator performance in Physcomitrella. This is not surprising, since both attributes are involved in the main function of terminators, namely terminating transcription and facilitating pre-mRNA processing into mature mRNAs through polyadenylation of the transcript’s 3’ end (Bentley, 2005; Mandel et al., 2008). Proper polyadenylation further influences mRNA export from the nucleus, transcript stability, and translation rates (Hoshino, 2012; Paek et al., 2015; Choe et al., 2018). However, PAS and poly(A) sites do not govern terminator performance alone. In Arabidopsis, stepwise deletion of PAS motifs from the AtHSP90t terminator did not reduce terminator performance (Andreou et al. 2021). In turn, 32 nucleotides at the 5’ end of AtHSP90t had a significant effect on reporter levels in *N. benthamiana*, even though this sequence does not contain any known motifs acting in polyadenylation (de Felippes et al., 2022). Combining the AtHSP90t 5’ with other terminators, similar to creating double terminators, further increased reporter levels. Respective transcripts did not differ in number or distribution of poly(A) sites or length of the poly(A) tail. Instead, a reduction in transcriptional read through and generation of small RNAs acting in transcript decay was observed.

In *N. benthamiana*, two terminators were identified and found to exceed reporter levels of the 35S terminator when tested in that same plant system (Diamos & Mason, 2018). These elements were identified by searching for stabilizing motifs in 3’UTRs to identify terminator candidates, based on Narsai et al. (2007), who identified stabilizing and destabilizing motifs in Arabidopsis 3’UTRs (Diamos & Mason, 2018). Interestingly, the list of stabilizing motifs includes the canonical PAS 5’-AATAAA-3’ as well as its reverse sequence 5’-AAATAA-3’, indicating RNA-binding *trans*-elements to be direction unspecific. Looking at other selection criteria used here, our results indicate that expression values did not have a large impact on terminator performance in Physcomitrella. At the same time, our results show the huge effect promoters have on gene expression as compared to terminators. Therefore, transcript abundance, which is the ratio of transcription rate to turnover rate, might be influenced to a larger extent by the promoter and therefore does not constitute an ideal criterion for terminator candidate selection. Likewise, the PCA demonstrated that terminator length did not exhibit a conclusive effect on terminator performance (**Fig. 5**). On the one hand, this observation fits with the findings by Lloyd et al. (2018), who showed that long 3’UTRs in Physcomitrella are not a positive predictor of NMD. On the other hand, we only characterized terminators in the range of 211 bp (35St) to 611 bp (Cab2t), which is still below the average length of 3’UTRs in Physcomitrella of 654 bp (**Fig. 1**). Consequently, our observation does not apply to Physcomitrella 3’UTRs in general but only to the length range tested here. In conclusion, of the selection criteria applied here for terminator candidate selection, the number of PAS and the number of poly(A) sites impact terminator performance notably, whereas terminator length and transcript abundance have a minor impact (**Fig. 5**). In addition, other factors such as the presence or absence of stabilizing or destabilizing motifs can act as good predictors for future terminator selection.

Since no single endogenous or heterologous terminator yielding higher than the 35S terminator could be identified, we analysed if combinations of two terminators would increase reporter gene levels. In Physcomitrella, the highest increases in signal strength by double terminators was achieved by the pairs Cab2t-petEt (ca. 1.6-fold). Of all the double terminators tested here, including pairs based on 35St and NOSt, as well as RPS6et and AtHSP90t, the pair Cab2t-petEt is the only combination which yielded more than the respective single terminators. As in *N. benthamiana* (Diamos & Mason, 2018), we also observed the effect of double terminators on gene expression to be dependent on relative positioning. This observation indicates that this effect is based on interactions between the respective terminators as opposed to additive effects. Whether heterologous or homologous pairs are preferable cannot be concluded here. The decrease in reporter signal levels for all combinations containing AtHSP90t is surprising. So far, AtHSP90t was combined with NtEUt, NOSt, NbHSPt, 35St, and itself, but for none of these pairs a negative effect of AtHSP90t was observed in *N. benthamiana* (Diamos & Mason, 2018; Yamamoto et al., 2018). Also, the pairs consisting of 35St & NOSt behave differently here as compared to other systems. Beyene et al. (2011) showed an increase in EYFP signal strength in transient particle bombardment assays in *N. tabacum* of up to 65-fold for 35St-NOSt in comparison to 35St alone. Diamos & Mason (2018) evaluated relative GFP and dsRed production via analysis of fluorescence levels visualized on UV-illuminated SDS-PAGE gels in *N. benthamiana* and found a change of >5 fold for 35St-NOSt compared to 35St, whereas other combinations reached >25-fold increases in reporter levels. One explanation for these discrepancies might be species-specific effects. However, 35St and NOSt are used as single terminators in a range of different plant species, including Physcomitrella. Since double terminators are generally functional in Physcomitrella and this effect is believed to be based on terminator-terminator interactions, it is surprising that 35St & NOSt double terminator pairs in Physcomitrella do not perform as well as in angiosperms. It is argued that double terminators lead to more stringent transcription termination and polyadenylation and less aberrant RNA, thereby reducing RDR6- mediated post translational gene silencing (PTGS) (Diamos & Mason, 2018). This argumentation is in line with findings in Arabidopsis where truncated and non-polyadenylated transcripts did not accumulate in wildtype plants but increased to higher levels in *rdr6* mutants (Luo & Chen, 2007). Further, usage of double terminators decreased mRNA 3’ read through and specific siRNA accumulation while increasing gene expression. It is argued that while RDR6-mediated PTGS might play a substantial role in explaining the positive effect of double terminators, it does not explain synergistic effects between terminators (Andreou et al., 2021). Instead, it is hypothesized that the positive effect of double terminators is also connected to promoter-terminator interactions. These interactions are believed to take place during gene looping when 5’ and 3’UTRs associate through bound members of the transcription machinery. A second terminator with additional and/or differing binding motifs for members of the polyadenylation complex could therefore interact more strongly with transcription factors bound to the 5’ end, resulting in more efficient gene looping and gene expression (Andreou et al., 2021). In Drosophila, a connection was found between the transcriptional start site (TSS) of a gene and the choice of poly(A)site, resulting in preferred TSS – poly(A) site pairs, which differed depending on the analysed tissue (Alfonso-Gonzalez et al., 2023). However, in Physcomitrella we observed the effect of double terminators on gene expression but found no clear evidence for promoter-terminator interactions.

Taken together, our data does not allow the deduction of any rules to predict how double terminators influence gene expression in Physcomitrella based on positioning and identity of the pair. However, observed effects do not behave additively but instead show positive and negative synergies. The mechanisms behind the effect of double terminators remains elusive. In general, double terminators can be an effective measure to increase molecular pharming yields which is why the mechanism behind double terminators should be elucidated further. For that purpose, methods to quantify transcriptional read through and PTGS and identify preferred poly(A) sites can be employed (Rosenthal et al., 2018; de Felippes et al., 2020; Shah et al., 2021). To investigate differences in transcriptional regulation between mosses and angiosperms, terminators with shown discrepancies between the two systems are of interest, such as 35St and NOSt pairs as well as AtHSP90t-based double terminators.

Besides species-specific effects, different reporter systems may be an explanation for discrepancies in characterization data of genetic elements. Most publications reporting large changes between different terminators and double terminators used a reporter system based on fluorescence proteins such as GFP and RFP (Beyene et al., 2011; Diamos & Mason, 2018; Rosenthal et al., 2018). While these results constitute promising reports for the utility of 3’UTRs and especially double terminators in boosting molecular pharming yields, other publications demonstrate a considerably smaller but nevertheless significant effect of terminators (Yamamoto et al., 2018; Tian et al., 2022; Chamness et al., 2023), in line with our findings in Physcomitrella. Yamamoto (2018) reported changes below two-fold in *N. benthamiana* for double terminators based on different combinations of AtHSP90t, NtEUt, and 35St using gel densitometry of Coomassie-stained SDS-PAGE gels. Using the Dual Luciferase reporter system in *N. benthamiana*, am less than two-fold difference between NOSt compared to NtEUt or 35St-NbACT7t-Rb7 was found (Chamness et al., 2023). The combination of 35St and NbACT7t with the matrix attachment region Rb7 was initially reported to yield a >60-fold increase in reporter levels compared to NOSt using a fluorophore-based reporter system (Diamos & Mason, 2018). In line with our findings, a maximum of 2.6-fold difference among 13 terminators characterized in both agroinfiltrated *N. benthamiana* leaves and BY2 cell culture was shown via the Dual Luciferase system (Tian et al., 2022). Further, the authors compared promoter characterization data obtained via Dual Luciferase assays to GFP reporter assays by verifying the data using RT qPCR. For the Dual Luciferase system, they found a correlation of 0.89 between firefly transcript and measured reporter values, whereas for GFP reporter assays a correlation of only 0.64 was found (Tian et al., 2022). Consequently, this indicates potential flaws in using fluorescence reporters for characterization of gene expression elements. A possible explanation might be the degree of autofluorescence emitted by plant cells and fluorescence signal interference with chlorophyll. In contrast, the Dual Luciferase system is based on chemiluminescence using unique substrates for both firefly and *Renilla* luciferases and is generally considered more sensitive than fluorescence-based assays (Naylor, 1999). However, the relatively short half-lives of firefly luciferase (2 – 3 hours) (Thompson et al., 1991; Ignowski & Schaffer, 2004) as compared to GFP (26 hours) (Corish & Tyler-Smith, 1999) might also explain the difference in the results obtained. Rosenthal et al. (2018) evaluated NtEUt using a GFP reporter, the GUS reporter system, and by quantifying Norwalk virus capsid protein via ELISA. The authors observed a 2.8-fold increase for NtEUt as compared to 35St using the GFP reporter. This difference decreased to around two-fold using the GUS reporter system and less than 1.5-fold when using ELISA, thereby nicely demonstrating discrepancies between the different reporter systems used in characterizing elements of gene expression. To conclude, characterization data for elements of gene expression can differ widely depending on the reporter system employed. This constitutes a problem regarding the comparability of published data, a concern that adds to the already existing complexity caused by species-specific effects.

In conclusion, we have described Physcomitrella 3’UTRs at the whole genome level, and identified and characterized 14 new endogenous terminator sequences. As a result, we obtained a selection of terminator sequences performing as well as established terminators 35St, AtHSP90t, and NOSt. Subsequent analysis of our terminator candidate selection criteria revealed polyadenylation-related motifs to be positive predictors of terminator performance. Double terminators also have the potential to increase gene expression levels in Physcomitrella, although no clear patterns in how double terminator composition affects outcome could be ascertained. Observed performance differences between single and double terminators in Physcomitrella and angiosperms may hint at evolutionary changes in the impact of terminators on gene expression.

## Statements & Declarations

### Author Contributions

P.A.N. designed research, performed experiments, analysed data, and wrote the manuscript. P.E. analysed data. S.W. performed experiments. J.P. and E.L.D. designed research. S.N.W.H. designed research, analysed data, and helped writing the manuscript. R.R. designed and supervised research, acquired funding, and wrote the manuscript. All authors read and approved the final version of the manuscript.

## Acknowledgements

We appreciate language editing by Anne Katrin Prowse.

## Data Availability

All terminator sequences identified in this study are deposited on GenBank.

| GenBank identifier | terminator |
| --- | --- |
| OR204709 | Cab1t |
| OR204710 | dehydt |
| OR204711 | petEt |
| OR204712 | Cab2t |
| OR204713 | Catalaset |
| OR204714 | PSIIt |
| OR204715 | UKt |
| OR204716 | rbcSt |
| OR204717 | EF1At |
| OR204718 | psaEt |
| OR204719 | RP-L18At |
| OR204720 | RP-S6et |
| OR204721 | RP-S18et |
| OR204722 | UK2t |

## Supplemental information

Supplementary file S1 **Vector maps**

Supplementary table S2 **List of primers**

Supplementary table S3 **List of vectors used in this study**

Supplementary table S4 **Terminator selection based on PEATmoss data**

Supplementary table S5 **Terminator selection based on microarray data**

Supplementary table S6 **Terminator selection based on GO term search**

Supplementary file S7 **Principal Component Analysis of reporter values and terminator attributes**

Supplementary tables S4, S5, and S6 are deposited on Zenodo (DOI: 10.5281/zenodo.8083448).

**Supplementary file S1 - Vector maps**

**Fig. S1.1.**
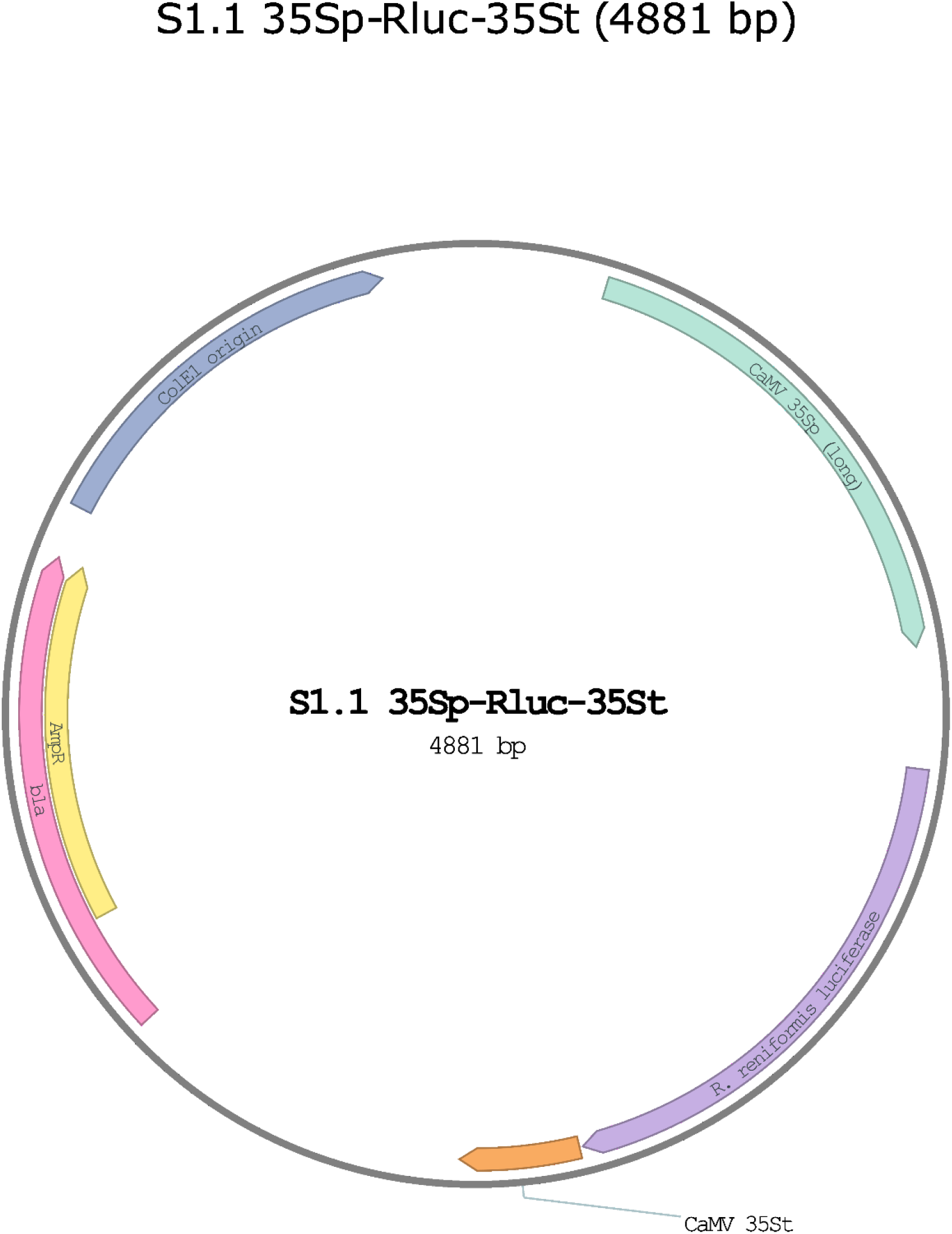
Vector map of 35Sp-Rluc-35St. *Renilla reniformis* luciferase-encoding vector used as a transformation control throughout experiments.

**Fig. S1.2.**
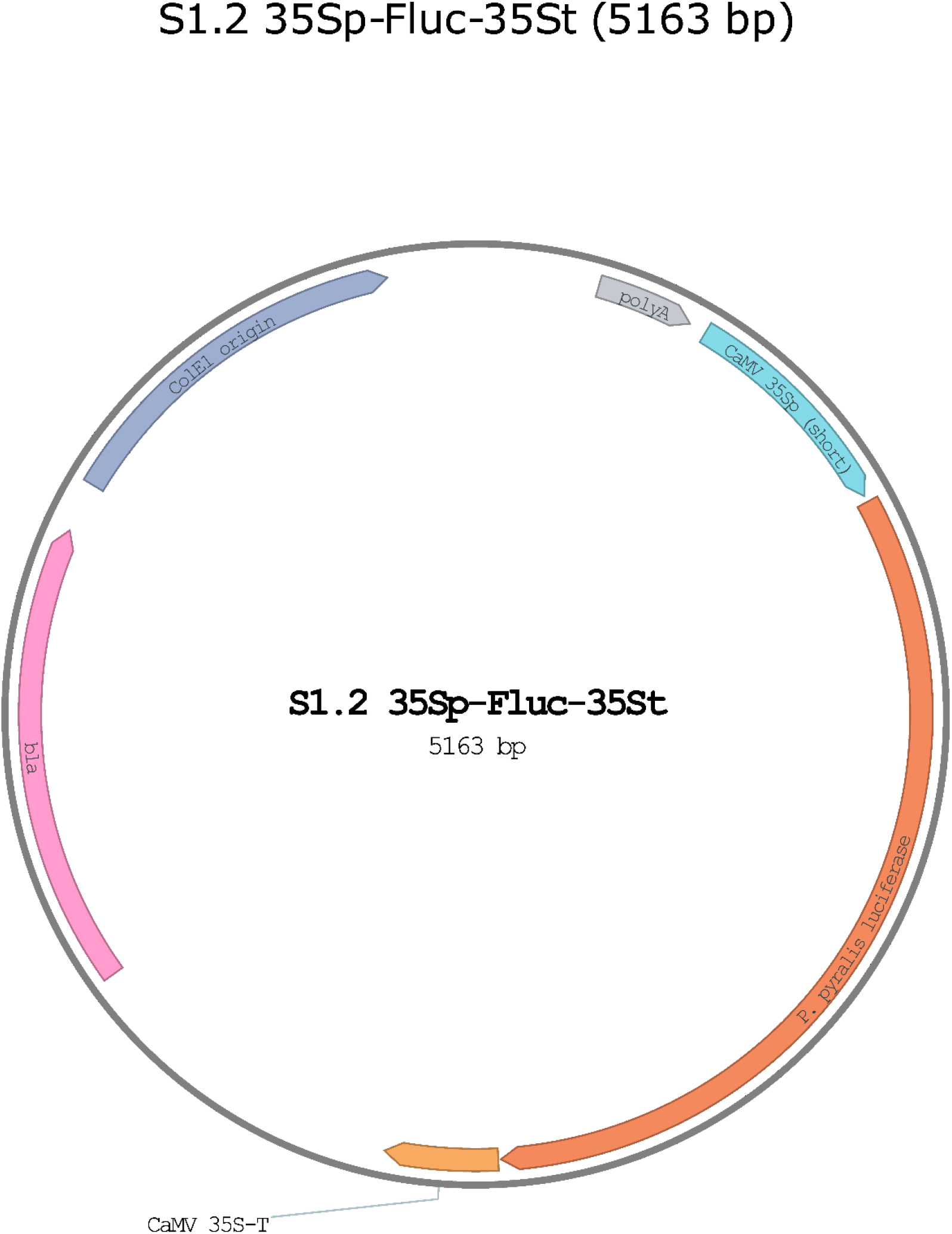
Vector map of 35Sp-Fluc-35St. *Photinus pyralis* firefly luciferase-encoding vector used as a reference throughout experiments and as a template for cloning of terminator testing constructs.

**Fig. S1.3.**
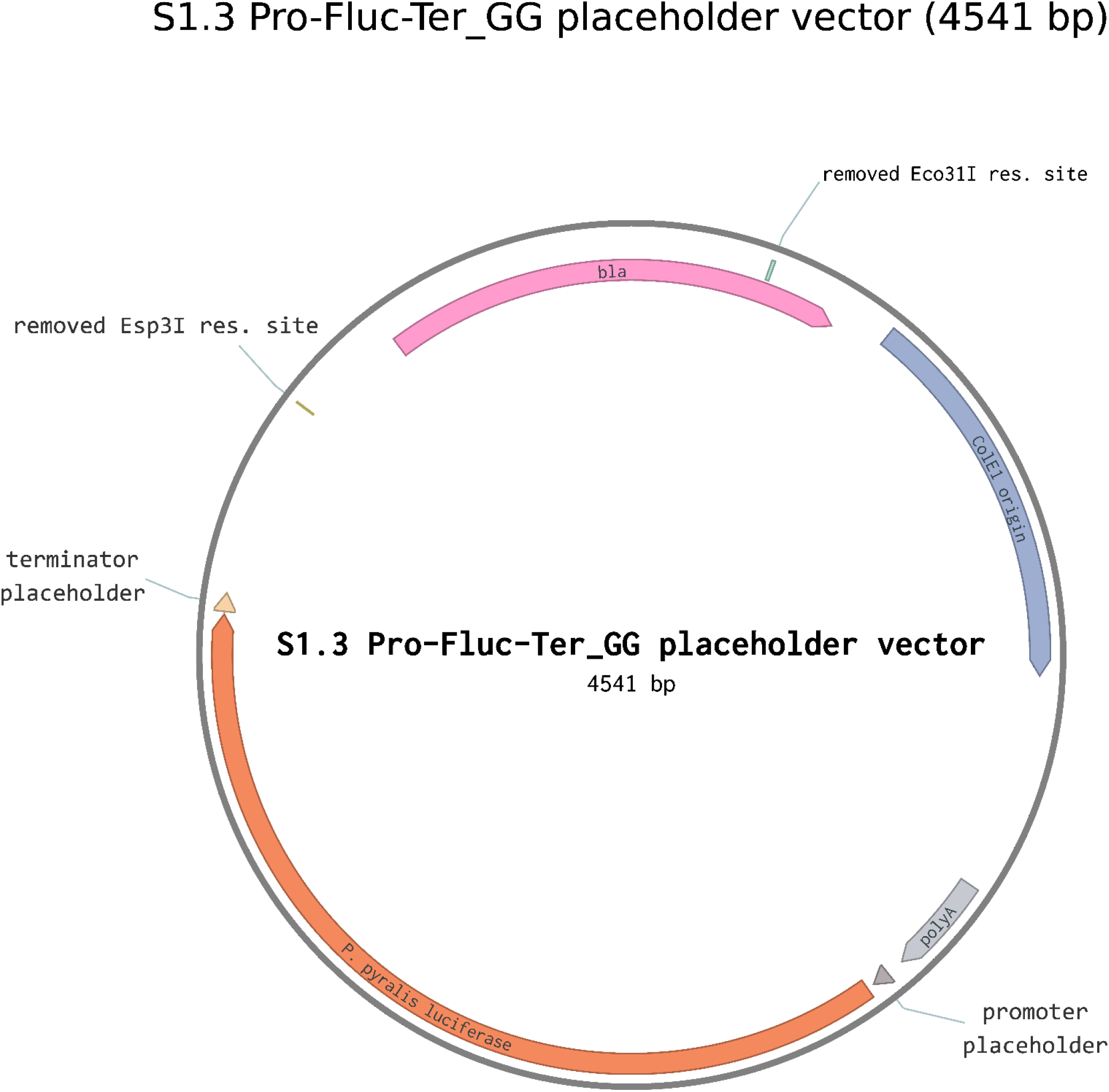
Vector map of Pro-Fluc-Ter_GG placeholder vector. *Photinus pyralis* firefly luciferase-encoding vector used as a template for cloning of terminator testing constructs via the Golden Gate-like cloning approach.

**Supplementary table S2.**
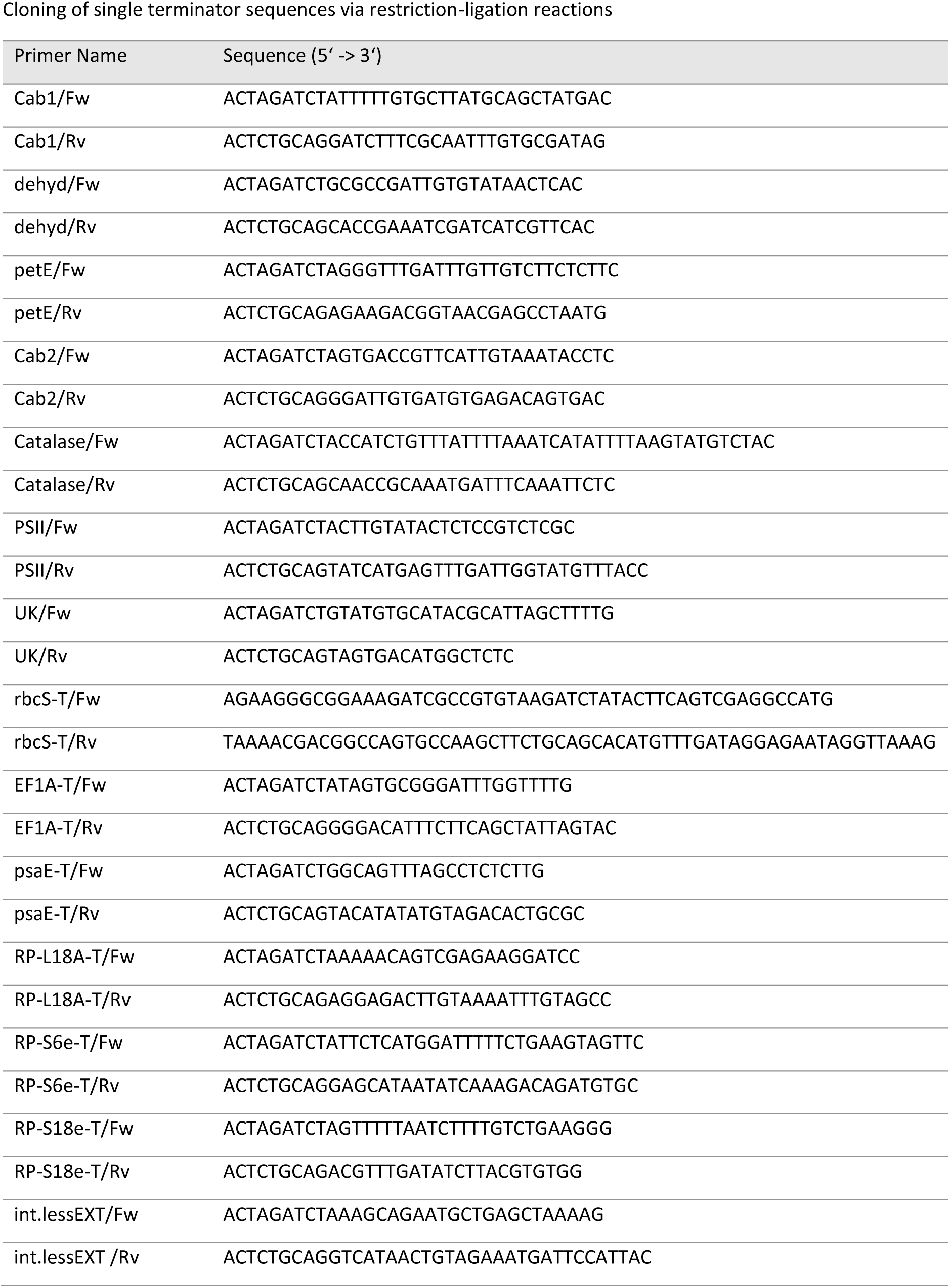
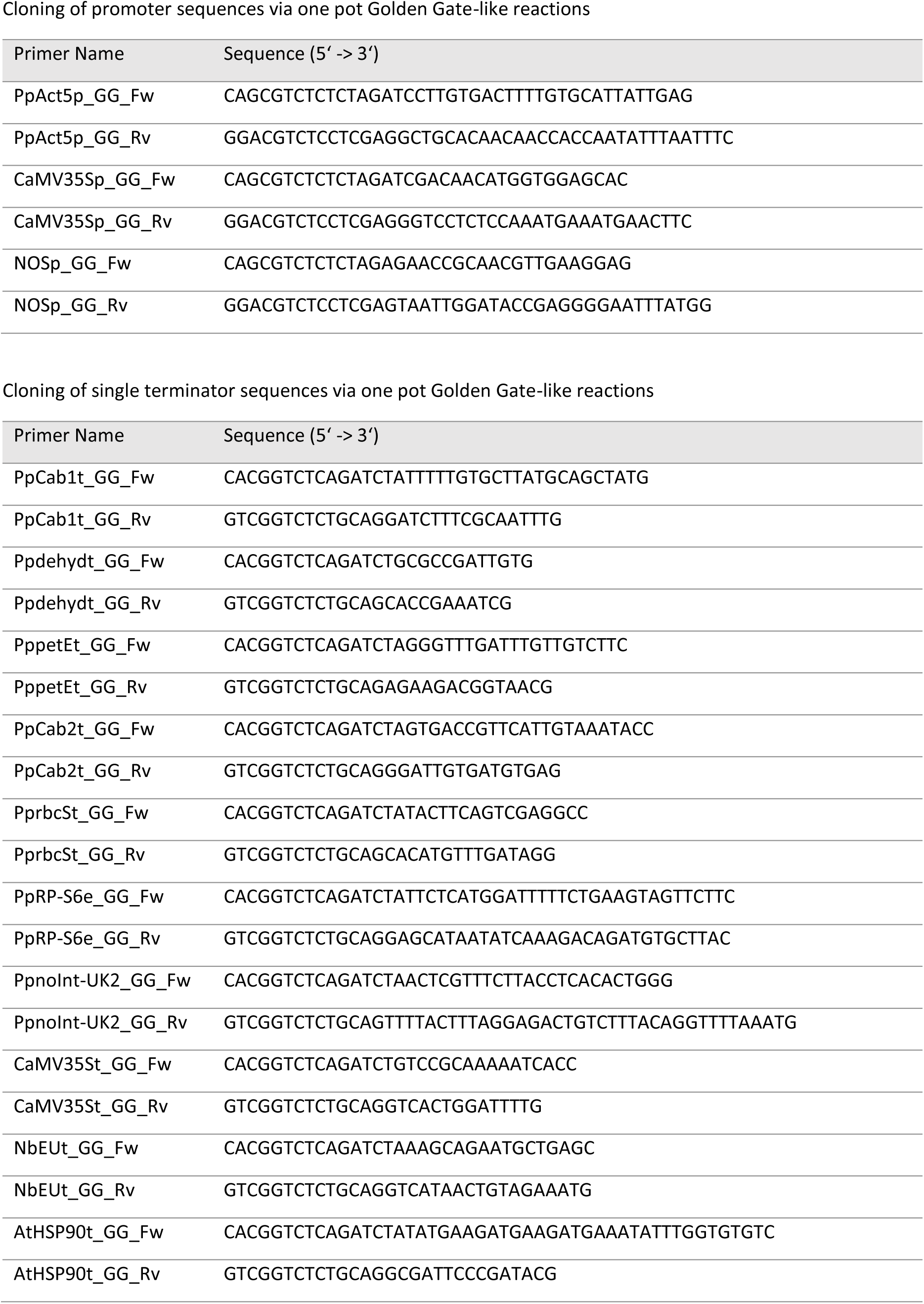
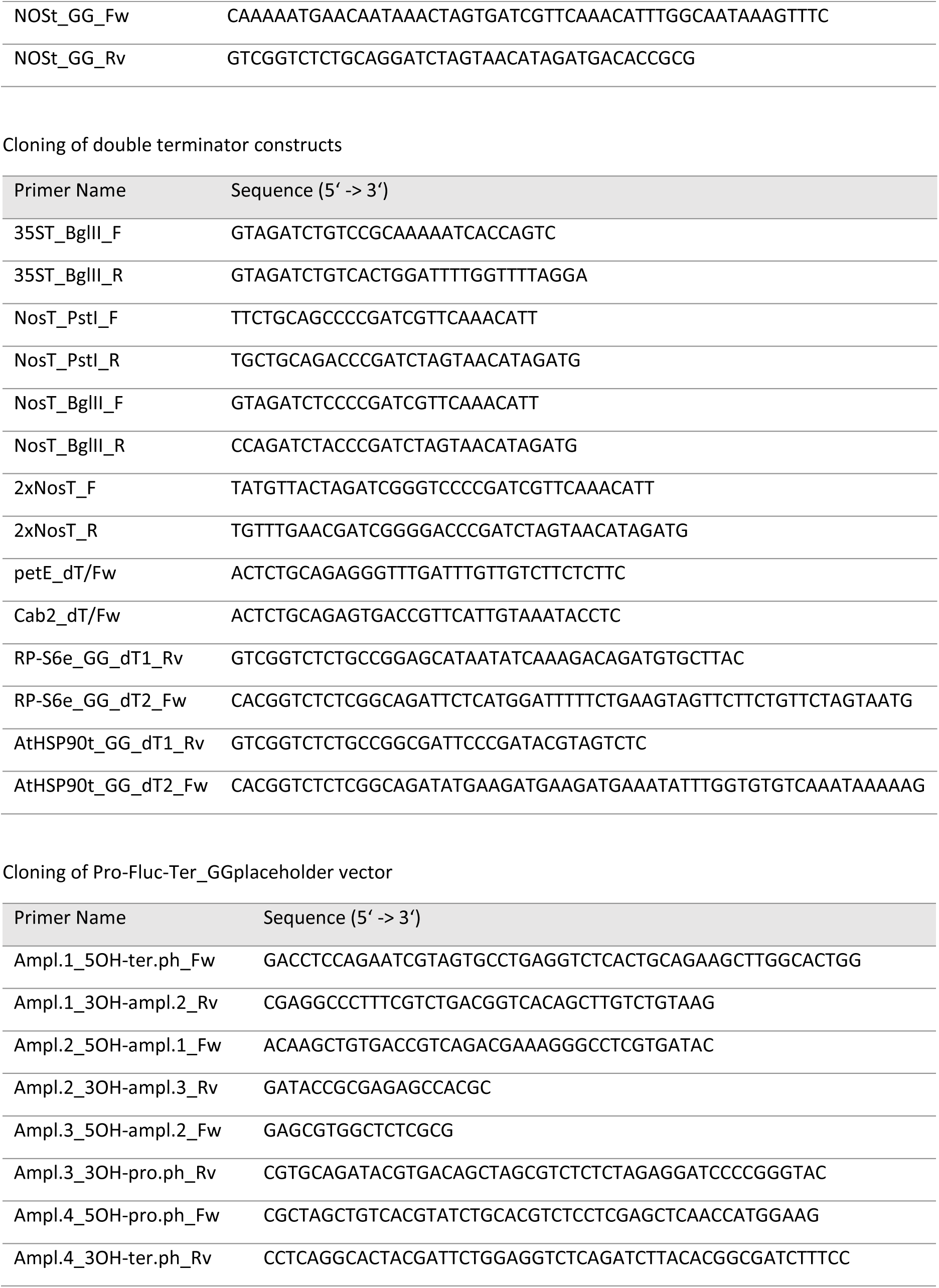
List of primers

**Supplementary table S3.**
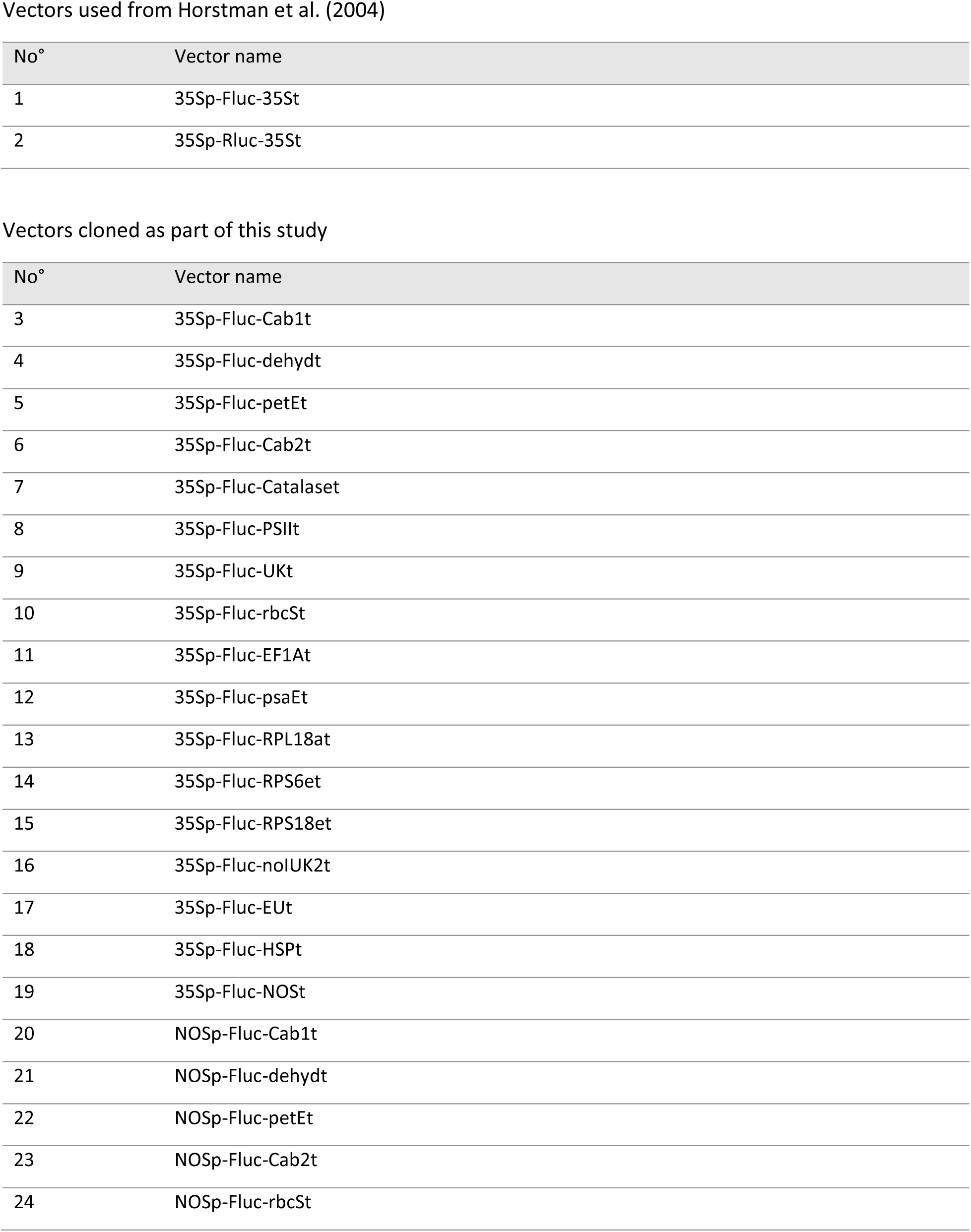
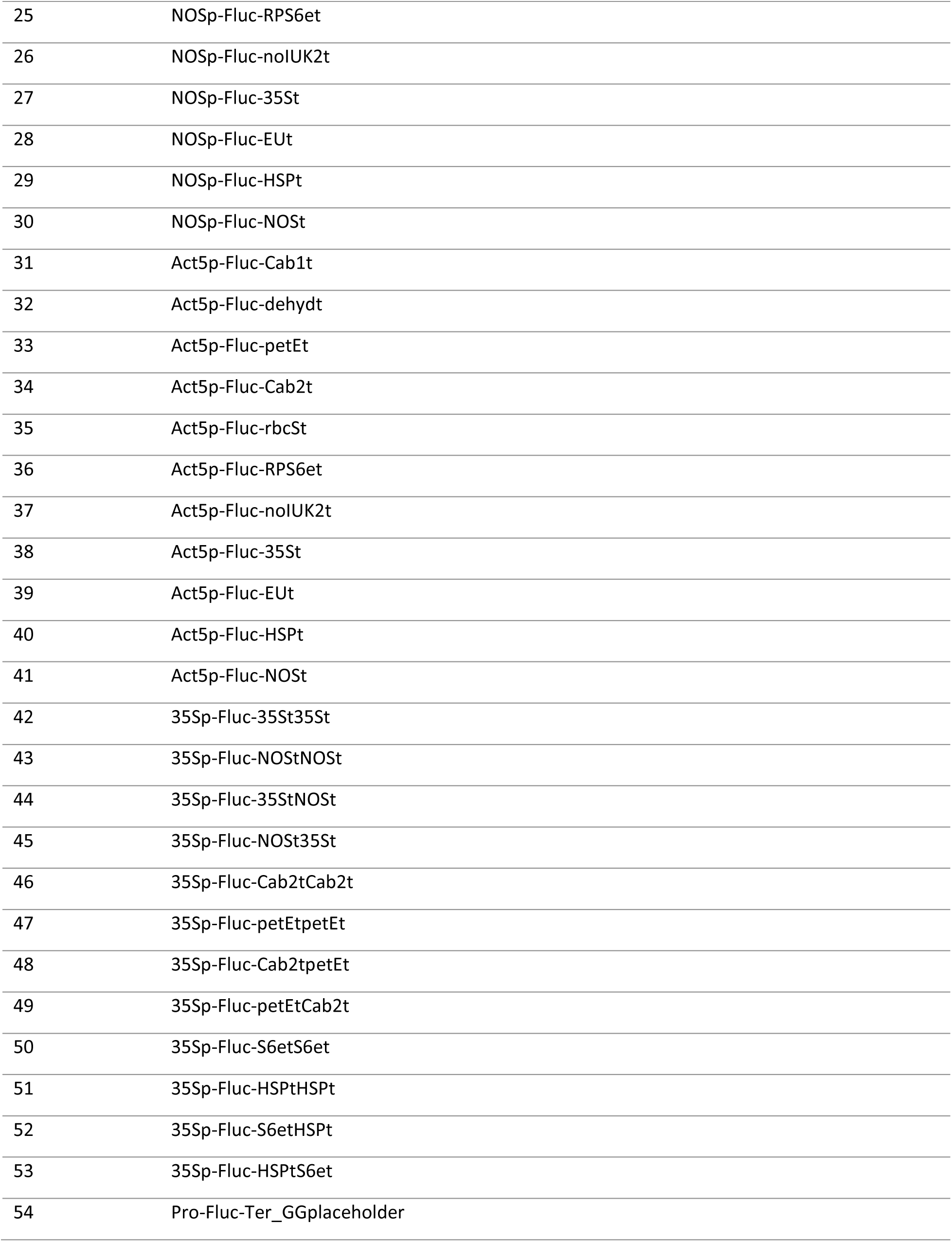
List of vectors used in this study

**Supplementary file S7.**
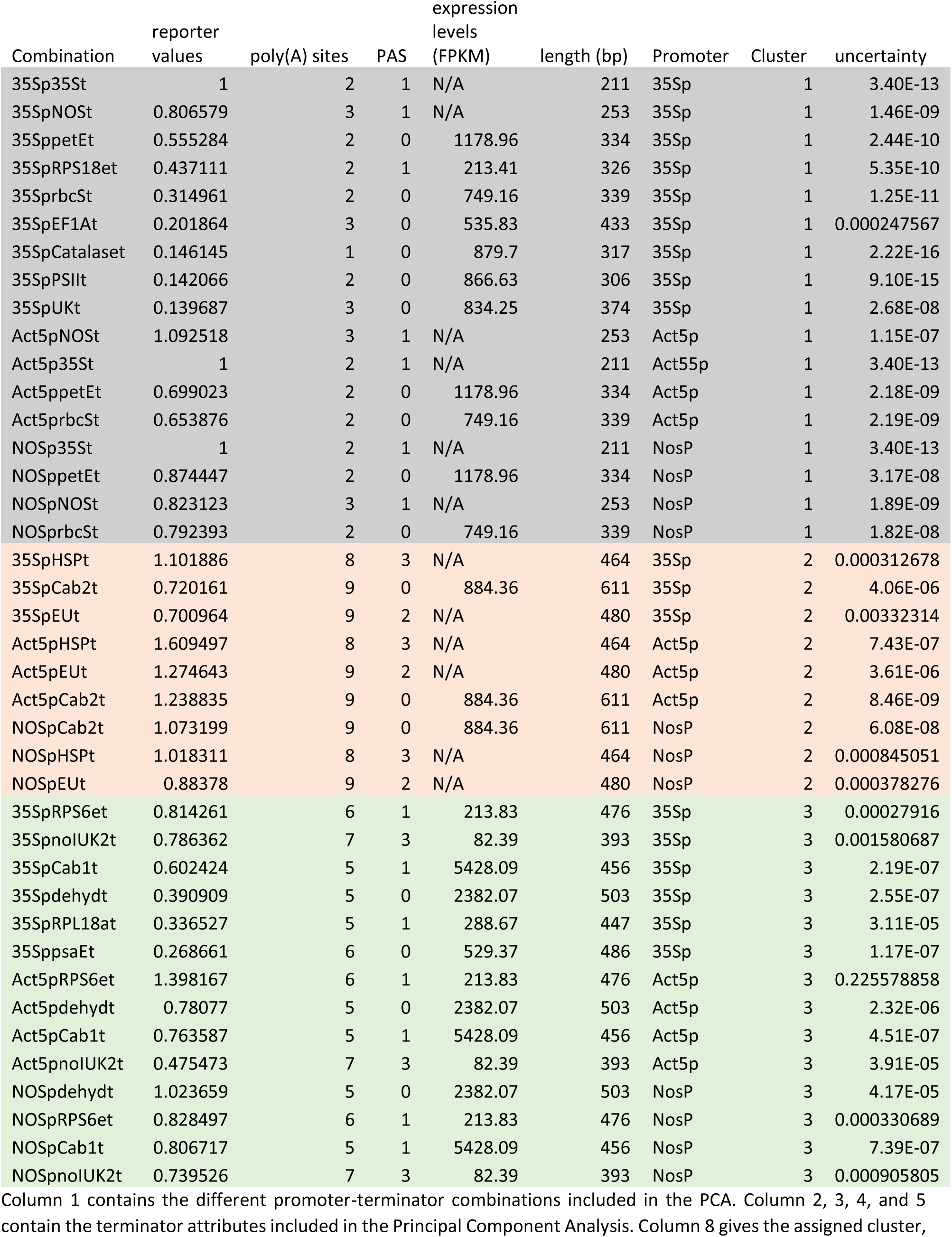
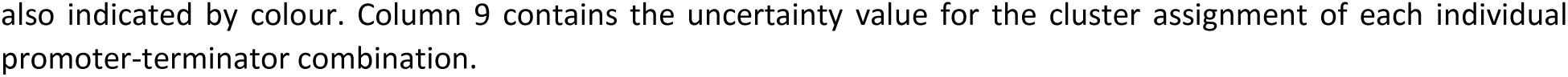
Principal Component Analysis of reporter values and terminator attributes.

